# Building a small brain with a simple stochastic generative model

**DOI:** 10.1101/2024.07.01.601562

**Authors:** Oren Richter, Elad Schneidman

## Abstract

The architectures of biological neural networks result from developmental processes shaped by genetically encoded rules, biophysical constraints, stochasticity, and learning. Understanding these processes is crucial for comprehending neural circuits’ structure and function. The ability to reconstruct neural circuits, and even entire nervous systems, at the neuron and synapse level, facilitates the study of the design principles of neural systems and their developmental plan. Here, we investigate the developing connectome of *C. elegans* using statistical generative models based on simple biological features: neuronal cell type, neuron birth time, cell body distance, reciprocity, and synaptic pruning. Our models accurately predict synapse existence, degree profiles of individual neurons, and statistics of small network motifs. Importantly, these models require a surprisingly small number of neuronal cell types, which we infer and characterize. We further show that to replicate the experimentally-observed developmental path, multiple developmental epochs are necessary. Validation of our model’s predictions of the synaptic connections using multiple reconstructions of adult worms suggests that our model identified the fundamental “backbone” of the connectivity graph. The accuracy of the generative statistical models we use here offers a general framework for studying how connectomes develop and the underlying principles of their design.

## Introduction

Biological neural networks develop according to “construction rules” that are genetically encoded [1–3], and are modulated and shaped by developmental noise [4], biophysical limits [5–7], and learning [8–12]. Uncovering the design principles that govern the diverse architectures of neural networks is fundamental to our understanding of the development, structure, and function of neural systems, and ultimately to our ability to interact with them, augment or fix them, and learn from them. Recent advances in reconstructing detailed connectivity maps, or *connectomes*, of networks of neurons at a resolution of individual cells and synapses [13–21] now make the quantitative study of these design principles tangible [22, 23]. Analyses of such networks have suggested that particular structural features are common, such as small-worldness and modularity [24], and the over-abundance of particular small network structures and the relative lack of others [25]. Following [26], a range of genetic, physical, and efficiency-driven features have been suggested to play important roles in shaping network topology and function [23, 27–29].

Probabilistic generative models offer a natural computational framework for modelling and studying connectomes [30, 31], as they allow us to explore which architectures and functions may arise from different construction rules and features and to conduct ablation experiments *in-silico*. Such models have been used to study brain wiring in different species, and could explain the observed “phase transition” in the relation between the number of synapses and the number of neurons during the growth of *C. elegans* [32]. They can also help identify neuronal cell types and predict their connectivity [33], and even generate accurate connectivity maps that also replicate neural activity, when simulated [23]. But, importantly, these models have not taken into account the developmental process itself, for the most part.

It is clear that reverse-engineering the design of complex networks and understanding them is difficult without modelling their development, even for relatively simple networks [34, 35]. Moreover, different network architectures can give rise to equivalent behaviors [36, 37], which makes the relations between structure and function hard to delineate by studying just the structural and functional endpoint of the developmental process. Thus, understanding the architectural and functional nature of connectomes may hinge on having a probabilistic generative model that can imitate the stochastic and gradual development of biological neural networks. Such models would offer us a way to explore both the fundamental building blocks, the construction plan, and the computation that these circuits may carry. They would also provide the means to explore circuit variability, robustness and the nature and causes of faulty circuits. Moreover, such developmental models may reveal different and hopefully simpler design principles of neural circuits.

We, therefore, present a novel family of generative developmental models, which rely on simple biological and physical features, for the study of the architecture and development of connectomes. We apply these models to the connectivity maps of the key part of the nervous system of *C. elegans*, as it is the only species to date for which multiple connectomes have been reconstructed at different developmental stages [16]. We show that models that we learn for one animal are highly accurate in predicting the detailed connectivity maps of held out parts of that connectome or of a full map of another individual - at the level of individual synapses, neuronal connectivity, and small network motifs. We further characterize the “developmental trajectories” our models predict and their tight similarity to the observed ones. Our models show that the connectome of *C. elegans* can be constructed accurately using a surprisingly small set of design rules. While these models offer a general framework that can be extended to other neural systems, our findings already suggest that a surprisingly small number of parameters govern the design of neural circuits.

## Results

To study the design of connectomes and the biological and physical features that shape their construction and architectures, we explore a family of computational models that generate connectomic maps, using different features. In particular, we ask how well they can recapitulate the experimentally measured connectomes of adults and during development.

### Generative developmental models for *C. elegans*

We begin by considering a simple family of generative models for the development of connectomes. These models construct neural circuits using three basic operations that are carried out iteratively, over discrete time steps (Fig. 1A): (1) Neurons are born at specific times and locations (as measured experimentally). (2) Synapses are formed independently of one another, with a probability that depends on the types of neurons they connect and the distance between them. Thus, the probability of forming a synapse from neuron *i* (which is of type *τ*_*k*_) to neuron *j* (of type *τ*_*l*_) in a given time bin is given by

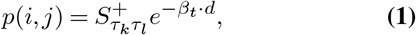

where 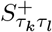 is the baseline probability to create a synaptic connection from a neuron of type *τ*_*k*_ to a neuron of type *τ*_*l*_, *d* is the distance between neurons, and *β*_*t*_ is a coefficient controlling the distance dependence, and its value is time-dependent to reflect the impact of the elongation of the worm during development (see Methods). (3) Individual synapses are removed via pruning with probability *S*^*−*^, in a manner independent of the types of neurons involved.

**Fig. 1.**
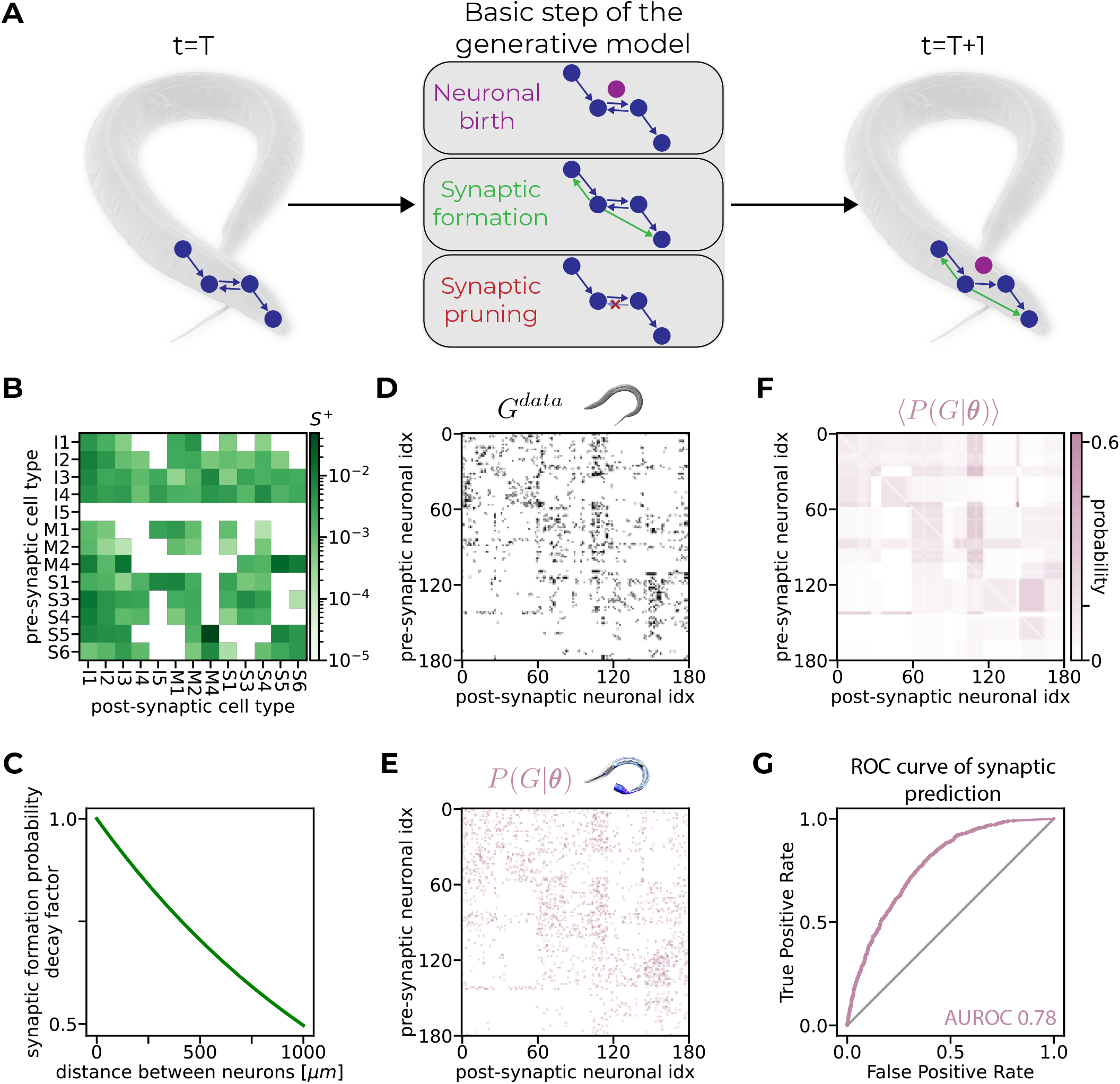
Generative models accurately predict the existence of individual synapses. **A**. An illustration of a basic step of the model, which comprises of the three basic operations iteratively used to construct the connectome: neuronal birth, synaptic formation, and synaptic pruning. **B**. A matrix of *S*^+^ values per each synaptic type after training (synaptic types are represented as a pair of pre- and post-synaptic neuronal cell types). **C**. The decay of the inferred baseline probability for synaptic formation as a function of the distance between neurons. **D**. The binarized connectivity matrix of the nerve ring of the adult worm used as test data (worm number 8 in [16]). Neurons in the matrix (and in the matrices in the following 2 panels) are grouped by type, and the ordering within a type is alphabetized according to their names. **E**. A connectivity matrix of a “synthetic worm” generated by a trained model. **F**. The average connectivity matrix of a trained model. Each entry in the matrix is the probability the model assigns for the existence of that synapse. Probabilities are exact (see Methods). **G**. The Receiver Operating Characteristic (ROC) curve of synaptic predictions. The probabilities in the matrix from **F** are used as predictions and the connectivity matrix from **D** as test data.

We use these models to study the connectome of the 180 neurons that compose the central part of the nervous system of *C. elegans*, known as the *nerve ring*. The detailed connectivity of the nerve ring has been reconstructed for several individual animals, at different time points during development [16]. Our model uses the binarized map of connectivity of the nerve ring of the adult worm. In this binary matrix, *G, G*_*ij*_ = 1 denotes the existence of at least one synapse from neuron *i* to neuron *j*, and *G*_*ij*_ = 0 denotes no connection. Fig. 1D shows this binarized connectome for one of the reconstructed adult worms from [16], which we denote as *G*^*data*^. Here, we use the common partition of the worm’s neurons into 15 functional types, from [15], of which 13 exist in the nerve ring (we shall revise this partition later). The birth times of individual neurons and their positions are taken from the measurements in [38] and [39], respectively.

For a given set of values of the parameters of our model 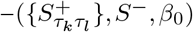,or ***θ*** for short – generating a connectome of the adult worm amounts to “running” it over time. Two neurons would be connected at the end point if they were wired together over development and were not pruned. This is akin to integrating over formation probabilities – *p*(*i, j*) from eq.1 – and the pruning probability – *S*^*−*^. We denote the probability that neurons *i* and *j* would be connected at the end point by *P* (*G*_*ij*_ = 1|***θ***). Therefore, the probability for a whole connectome, *G*^*′*^, is given by

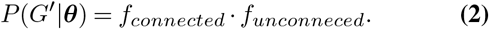

Namely, *P* (*G*^*′*^|***θ***) is the product of the probabilities of all the pairs of neurons that have synaptic connections,

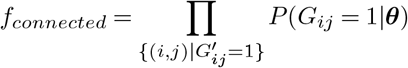

and the product over all “non-existing” synapses, given by

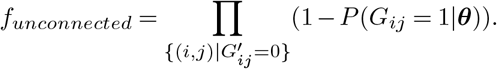

We then seek the set of ***θ*** that would maximize the likelihood of the data, *P* (*G*^*data*^|***θ***), while retaining the experimentally measured average number of synapses between each pair of neuronal cell types (see Methods and Fig. 1B,C).

### Generative developmental models accurately predict individual synapses in the adult worm

We use the model we have learned, *P* (*G*|***θ***), to synthesize connectomes; one such “draw” from the model is shown in Fig. 1E, and the average synthesized connectome is shown in Fig. 1F. We quantify the performance of the model by the accuracy with which it correctly predicts the existence of individual synapses in the measured connectome, or the lack thereof. Since the model assigns a probability for the existence of each synaptic connection, we use the Receiver Operating Characteristic (ROC) curve over all the synapses to characterize the synaptic maps (as done in [23]), and summarize the performance of the model by the area under the ROC curve (AUROC). Since the connectomes of the nerve rings of two adult worms have been reconstructed in [16], we used one of the worms for training the model and the other one as test data.

Our model achieves an AUROC value of 0.78 using 13 neuronal cell types on test data (Fig. 1G). Notably, this value is similar to the accuracy of probabilistic generative models of the adult worm connectome that do not rely on developmental processes (and are therefore less constrained) [23].

We also asked how sensitive is our model to the exact timings of neuronal birth, and found it to be robust to a significant level of “temporal noise”. Specifically, differences of up to 50% in the birth times of individual neurons had a very small effect on the accuracy of prediction of individual synapses or the wiring diagram as a whole (Fig. S 1), consistent with [28].

In the next section, we show that we can achieve an even higher level of accuracy with fewer parameters. This implies that a much more compact set of neuronal cell types may shape connectivity.

### A small number of inferred neuronal types is sufficient for building an even more accurate adult connectome

To assess the contribution of different structural and biophysical features to predicting connectivity, we evaluated the performance of model variants that depend on subsets of the features of *P* (*G*|***θ***) (Fig. 2A): (1) A model based solely on the birth times of neurons; (2) A model based solely on the distances between neurons (see Methods). Both the birth-time- and distance-based models perform above chance in predicting the existence of individual synapses (Fig. 2B). We then evaluated the performance of *P* (*G*|***θ***) as a function of the number of neuronal cell types it relied on, using either a single neuronal cell type, 3 neuronal cell types (sensory neurons, motor neurons, and interneurons), or the 13 neuronal cell type assignments taken from [15]. We found that the performance increased with the number of neuronal cell types (Fig. 2C), raising the questions of how many cell types would saturate the accuracy of the model, what would these types be, and how can they be identified.

**Fig. 2.**
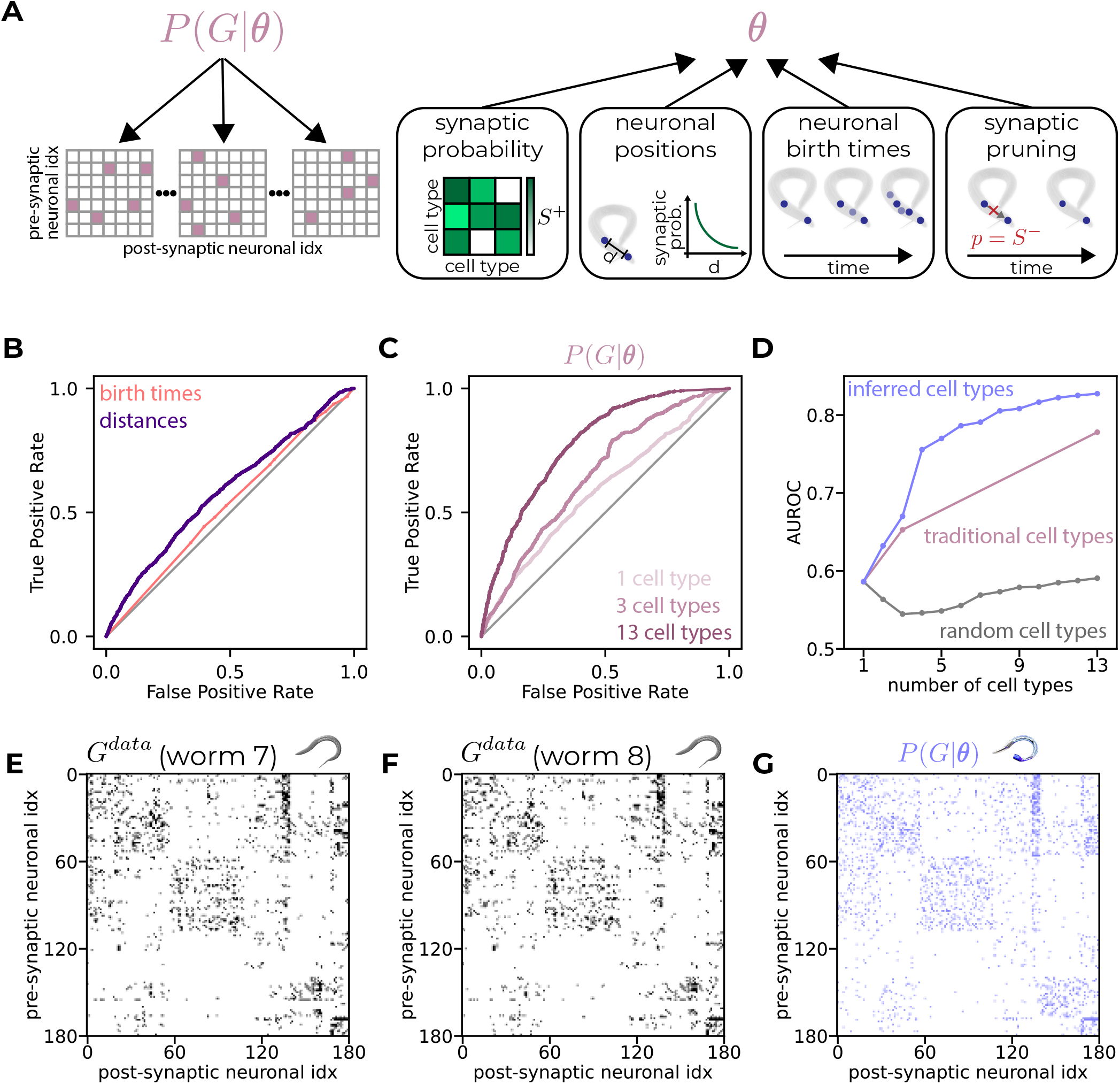
A small number of neuronal cell types is sufficient to generate accurate connectomes. **A**. An illustration of the different features used to construct the variants of the model. Left - *P* (*G*|***θ***) generates synthetic connectivity maps. Right - ***θ*** is comprised of baseline synaptic formation probabilities, one for each pair of neuronal cell types (the *S*^+^ matrix); positions of neurons in the body of the worm and the decay of synaptic formation probability with distance; neuronal birth times; and synaptic pruning, which is done with probability *S*^*−*^. The number of neuronal cell types of models may vary. **B**. ROC curves of 2 model variants using one neuronal cell type (all neurons have the same type). **C**. ROC curves of *P* (*G*|***θ***) when using a single cell type, 3 cell types (sensory, motor and interneurons) or 13 cell types following [15]. **D**. AUROC of *P* (*G*|***θ***) versus the number of neuronal cell types used, when using random types, traditional cell types from [15] or inferred cell types. **E-F**. Binarized connectivity matrices of real connectomes (worms 7 and 8 from [16] in panels **E** and **F**, respectively). Neurons in the matrix (and in the matrix in the next panel) are grouped by type and the ordering within a type is alphabetical according to their names. **G**. The connectivity matrix of a synthetic connectome generated by a model that uses 8 inferred cell types.

We, therefore, asked whether there is a different classification of the neurons into cell types that would give a more accurate model than the ones based on the traditional classification of neurons into types defined in [15]. We used a simple heuristic agglomerative clustering of the neurons into new types that collects together neurons so that a model based on these new types would be most similar to the data (see Methods). Surprisingly, we found that models based on these inferred types are much more accurate and rely on far fewer types (Fig. 2E-G): the performance of a model that uses only 4 inferred cell types was similar to that of a model that used the “traditional” 13 types from [15] (Fig. 2D). As the AUROC of the models improved rapidly up to 10 inferred cell types, and far less for more than 25 types (Fig. S 2A), we focus henceforth on models that rely on 8 inferred cell types, where the AUROC value crosses 0.8. We further validated that this choice does not result in over-fitting (SI and Fig. S 2B). We explore the nature of these types below (and note that since our classification into types relied on a heuristic clustering, there may be an even more compact set of types that would perform even better).

### Generative models based on small number of inferred types of neurons recapitulate neuronal and sub--network features of the measured connectomes

We next assessed the performance of *P* (*G*|***θ***) beyond individual synapses, and quantified the accuracy of predicting connectivity features of single neurons, as well as small sub-circuit structures. Fig. 3A shows the similarity of the in-degree and out-degree distributions predicted by *P* (*G*|***θ***) and the experimental data, as the observed degree frequencies lie within 2 standard deviations (std) from the mean of the model’s predictions, for almost all values.

**Fig. 3.**
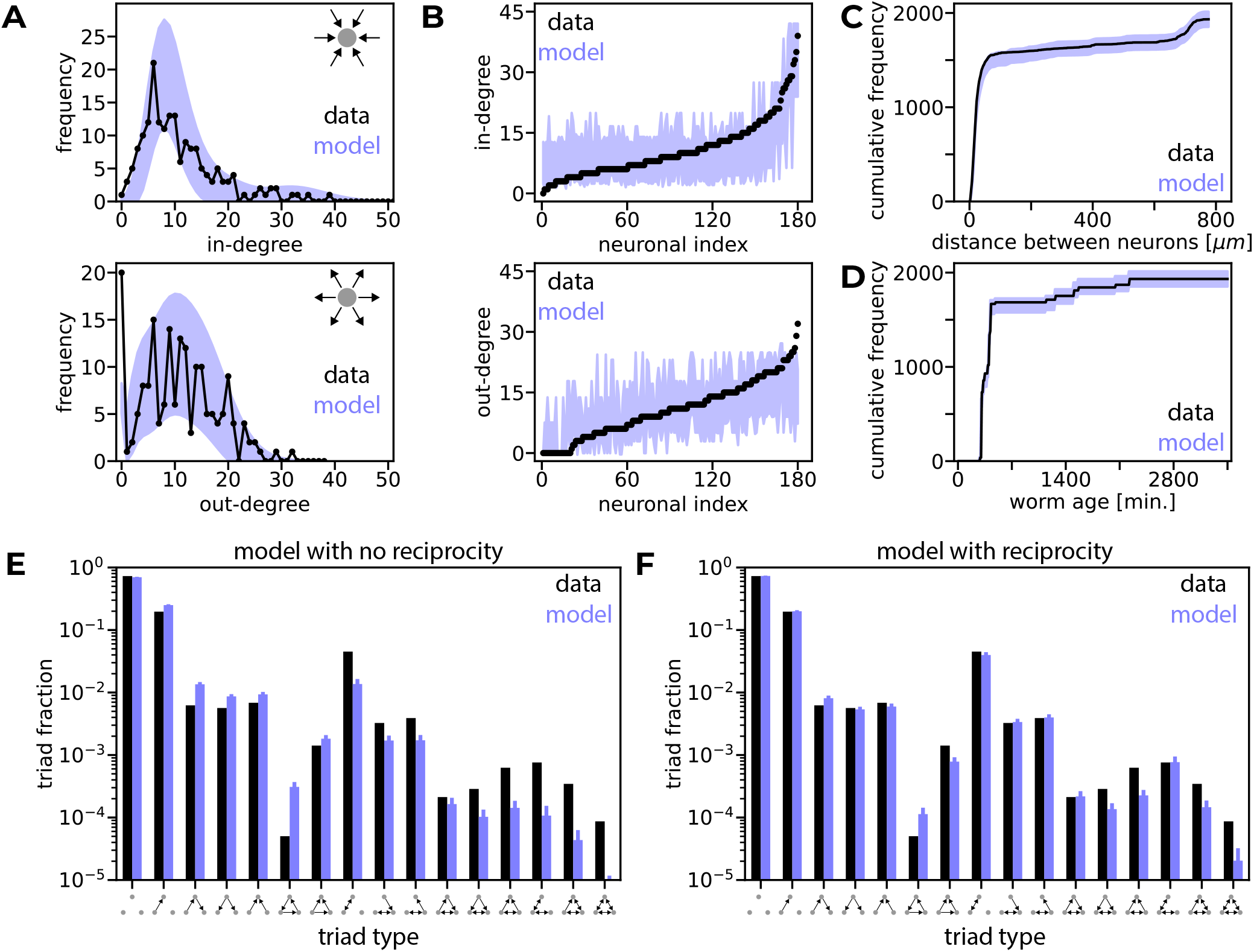
Models reproduce neuronal and network properties. **A**. Distributions of in-degrees (top) and out-degrees (bottom) of the data and the model. Shaded areas represent 2-std around the mean degree distribution of the model. Calculations are exact (see Methods). **B**. Degrees (in-degrees on top and out-degrees on bottom) of individual neurons in the data and the model. Neurons are sorted in an ascending order of their degrees in the data. Shaded areas represent 2-std around the mean degree in the model. Calculations are exact (see Methods). **C**. A cumulative histogram of the distances between connected pairs of neurons in the model and the data. Shaded areas represent 2-std around the mean distribution of distances in the model. Calculations are exact (see Methods). **D**. A cumulative histogram of the maximal birth times of pairs of connected neurons in the model and the data. Shaded areas represent 2-std around the mean distribution of maximal birth times in the model. Calculations are exact (see Methods). **E**. The triplet motif distribution of the data and the mean distribution of the model. Error-bars show 2-std. Calculations are numerical (see Methods). **F**. The triplet motif distribution of the data and the mean distribution of a model with statistically dependent reciprocal synapses (*γ* = 7) and 8 neuronal cell types. Error-bars show 2 std. Calculations are numerical (see Methods.)

The distribution of the number of synaptic connections of specific neurons in *C. elegans* has resulted in the designation of some neurons as “hub neurons”, which were suggested to play an important role in information processing in the worm [40]. We, therefore, compared the in-degree and out-degree values of individual neurons as well. Here, too, we found that the model predictions agree with the data, as most of the degrees lie within 2-std from the model’s mean (Fig. 3B).

We found that *P* (*G*|***θ***) reproduces additional spatial and temporal features of the synaptic map. Fig. 3C shows that the model accurately predicts the histogram of physical distances between connected neurons. We also compared the times at which synaptic connections may form, using the first time point at which both neurons of a synapse exist, and found that this synaptic feature is reproduced as well (Fig. 3D).

We emphasize that the accuracy of *P* (*G*|***θ***) in recapitulating characteristics of neuronal connectivity does not result from explicit features shaping the neuronal degrees in the model. Moreover, while our model includes dependency on the distances between neurons and on neuronal birth times, it was not trained to recover the histograms of these values over the connected neurons.

We then compared the distribution of connectivity patterns among all triplets of neurons in the measured connectome with the predictions of *P* (*G*|***θ***). We found that our model recovered the relative frequencies of most of these triads, but did not capture their exact values (Fig. 3E). Specifically, the model underestimated the frequency of triads that contained reciprocal connections between neurons, and over-estimated ones that did not. This suggests that our model lacks some design features that shape the sub-circuit frequencies, which we turned to explore next.

### Models’ inaccuracies reveal new design features of the adult connectome

The inaccuracy of our current model in predicting triplet connectivity patterns implies that synaptic connections are not independently formed. This notion is corroborated by the fact that the reciprocity of synaptic connections in the experimental data is much larger than that projected by the Erdős-Rényi (ER) model [41] for a network with the same synaptic density. In ER networks, the average reciprocity is given by the square of the density of synaptic connections, which in the case of the nerve ring of *C. elegans* would be *∼* 0.0036, or *∼* 4.5 times smaller than the observed reciprocity in the data. Our current model, *P* (*G*|***θ***) – which relies on 8 inferred cell types, the distance between neurons, and the birth times of the neurons – has a reciprocity value which is *∼* 1.7 times that of the ER model (Fig. S 3A). Notably, using more neuronal cell types only mildly improves the reciprocity prediction.

We, therefore, explored an augmented version of the model, in which the probability for the formation of a synapse from neuron *i* to neuron *j* increases if a synapse from *j* to *i* already exists. This probability is given by min 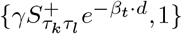 where *γ* is an additional reciprocity parameter (see Methods). The reciprocity of this new model as a function of *γ* (using the same 8 neuronal cell types as before), shown in Fig. S 3B, demonstrates that a model with *γ* = 7 results in reciprocity that is close to the observed value (i.e., the observed value is within the variance of the model’s outcomes). Importantly, the reciprocity parameter fixes most of the estimates of the triads’ distribution (Fig. 3F) without changing the AUROC values for predicting individual synapses or the in-degree and out-degree distributions. To quantify the improvement, we calculated the Jensen-Shannon divergence between the observed distribution and the average distribution of the model; the divergence decreased from 0.01135 to 0.00045 bits. Thus, the failure of the original version of the model unveiled the tendency of reciprocal synapses to be formed as a missing design feature.

### The inferred neuronal cell types of the model have clear biological interpretations

The highly accurate performance of the updated generative model *P* (*G*|***θ***) – that now relies on inferred neuronal cell types, the distance between neurons, neuronal birth times, and reciprocity – raises the question of the nature of these inferred cell types. Fig.4A shows the overlap between the traditional cell types of [15] and our inferred cell types, reflecting that the inferred cell types are a rehashed and more compact classification. We further show that our inferred types designate clear biophysical features, as they reflect neuronal relative locations and their birth times (Fig. 4B). Unlike the cell types from [15], our inferred neuronal cell types do carry information regarding the in-degree and out-degree values of individual neurons (Fig. 4D). Moreover, our inferred types group together hub neurons: Fig. 4E shows the relation between the number of neurons assigned to each inferred type and its directional hub value (given by max{*⟨d*_*in*_*⟩,⟨d*_*out*_*⟩*} of that type, where *d* stands for degree). Our results suggest that the connectivity profile itself may be a biological feature of neurons, consistent with the finding that the similarity of gene expression profiles of hub neurons in *C. elegans* cannot be explained by other biological features that they share [42].

**Fig. 4.**
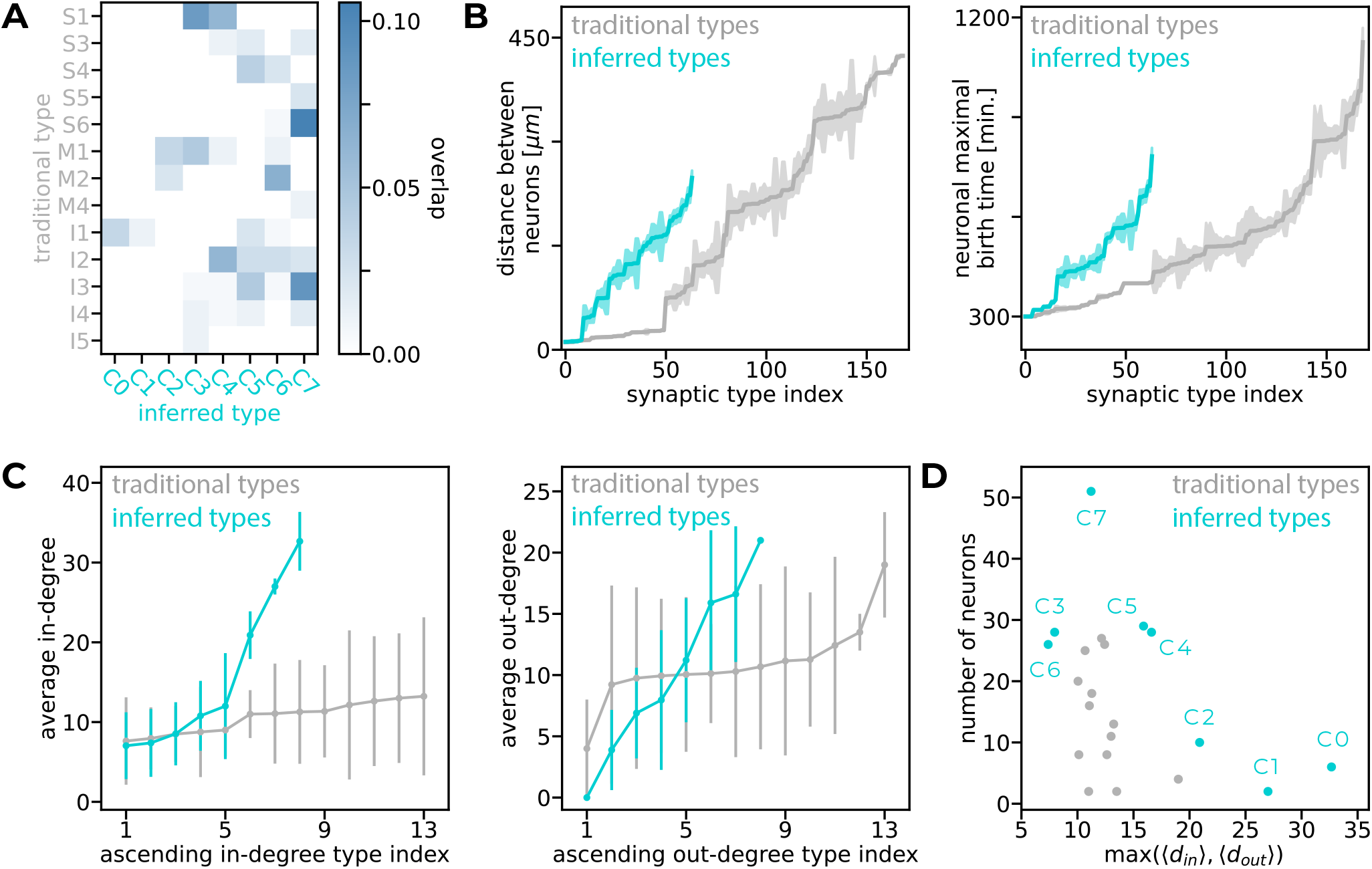
Biological interpretation of the inferred neuronal cell types. **A**. The mixture matrix of the 13 traditional cell types from [15] (vertical axis) and the 8 inferred cell types (horizontal axis). Each entry of the matrix reflects the fraction of neurons that have the traditional type (rows) and the inferred type corresponding to it (columns). Thus, the sum of all the entries in the matrix is 1. **B**. Left - average distance between neurons versus the index of their potential synaptic type (defined as a pair of neuronal cell types). Synaptic type indices are ordered such that the average distance is monotonic. Right - average maximal birth time of pairs of neurons versus the index of their potential synaptic type. Synaptic type indices are ordered such that the average maximal birth time is monotonic. Shaded areas show the standard error of the mean (SEM). **C**. Average degrees (left – in-degrees, right – out-degrees) of neuronal cell types in the data. Types are sorted in an ascending order (the indexing is different for in- and out-degrees). Error-bars show std. **D**. Type size (the number of neurons assigned to that type) versus the directional hub value, given by max{⟨*d*_*in*_⟩, ⟨*d*_*out*_⟩} of that type, where *d* stands for degree.

### Further compression of the generative model shows there is no fine-tuning of the parameters for its accurate and robust performance

Our developmental model based on inferred neuronal types depends on far fewer parameters than models that rely on the 13 types from [15], or the 118 neuronal classes that were defined based on anatomy and molecular markers [43, 44]. This compression raises the computational and biological question of the minimal model needed to construct the worm’s nervous system.

We found that we can achieve a highly accurate model by the quantization of 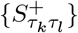(see Methods, Fig. 5A,B). The model’s performance increased rapidly with the number of values and saturated already for 14 parameters (namely, a compression of almost 5 fold of the full model’s size). Specifically, a model with just 9 values achieved an AUROC value of 0.77, which is 95% of the accuracy of the full model with 8 types (AUROC of 0.81), over the full connectivity map with *∼* 30*K* potential synapses (Fig. 5C). Fig. 5D shows that the compression is robust to the number of cell types chosen: increasing the number of types changed the size of compressed models only mildly, while the size of the full model grew fast. These results reflect the high robustness of *S*^+^ in terms of the numerical accuracy of its individual values. We conclude that a surprisingly small number of parameters is sufficient to construct the connectome of the worm to a high degree of accuracy, and these parameters are not fine-tuned.

**Fig. 5.**
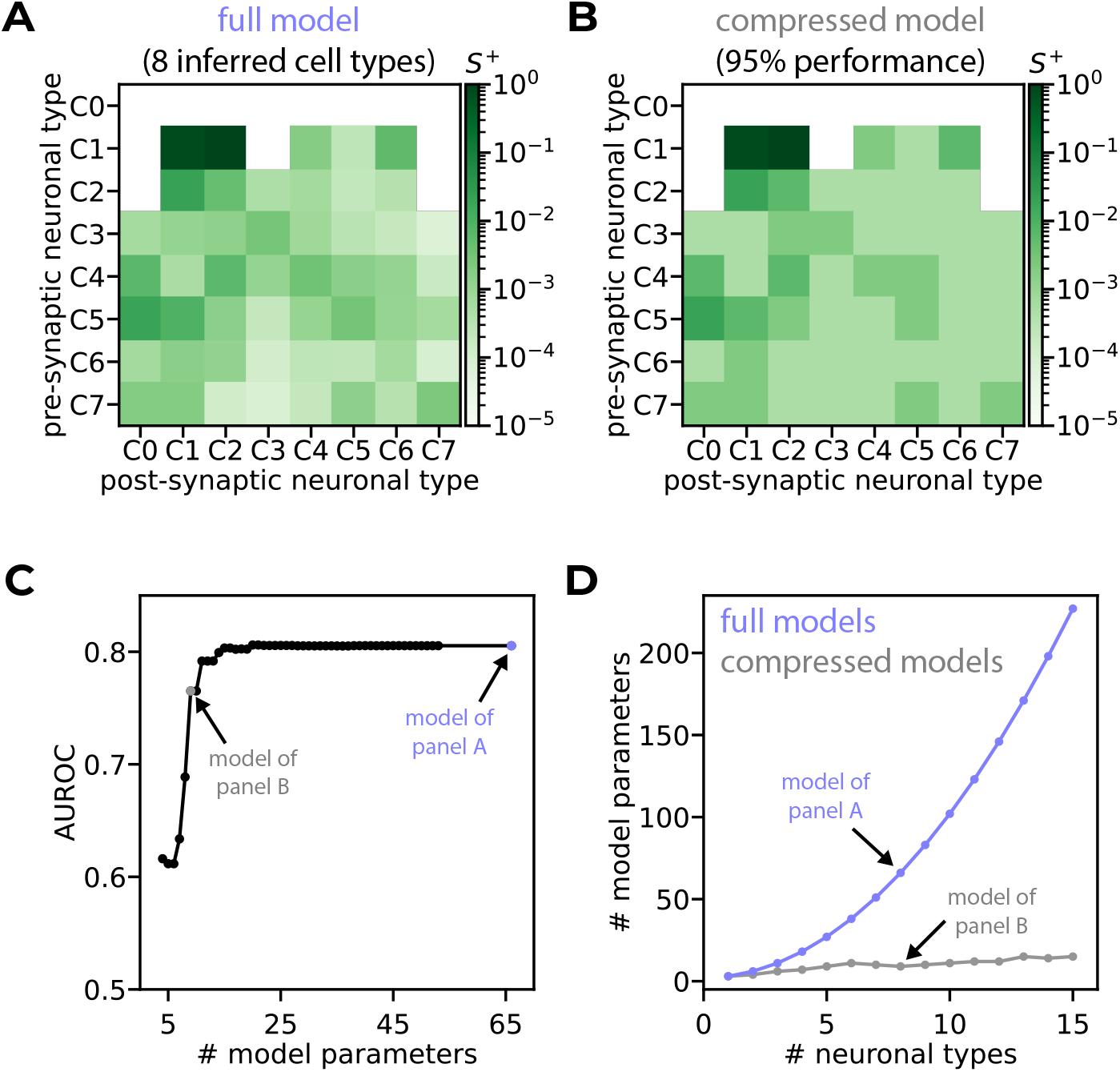
Highly compressed models preserve accuracy. **A**. The matrix of *S*^+^ values of a model that uses 8 inferred cell types. **B**. A quantization of the matrix from **A** that conserves 95% of the AUROC value using 7 distinct values. **C**. The AUROC of models based on 8 inferred cell types that were compressed to different sizes versus the number of model parameters. **D**. The number of model parameters versus the number of neuronal cell types for full models and for compressed models that conserve 95% of the AUROC value.

### Multiple developmental epochs are necessary to recapitulate the measured developmental path

Given the accuracy of our generative model for the adult worm, we turned to study the development of the connectome, using the reconstructions of the nerve ring of 8 different nematodes from [16]. Given that for young larvae there is only a single reconstruction available per age, to learn our models and validate them, we randomly split the total set of potential synapses of the adult worm into two halves. One half was used as training data and the other as test data (see Methods) for all developmental stages. We compared the accuracy of *P* (*G*|***θ***), which relies on a single set of parameters for the whole developmental process, with models that use three developmental epochs to predict the empirical connectome (Fig. 6A, Fig. S 4). The values of 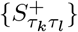in the latter model may differ between epochs, which were defined as the time windows between the developmental stages of the reconstructed connectomes of two different young larvae and the adult developmental stage. Models were trained sequentially, such that we found ensembles of 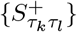 (one ensemble per epoch) that maximize the likelihood of the adult worm, while conserving the number of synapses of each type in three of the worms across development (see Fig. S 5, Fig. S 6, Methods). The performances of the single epoch model and the multiple epochs model in predicting the connectome of the adult nematode were surprisingly similar (Fig. 6B). However, this does not necessarily mean that the models have similar performance during development, as they may reach the same endpoint even if they follow divergent trajectories in the space of networks (Fig. 6C). Indeed, the model based on a single epoch and the one based on multiple epochs follow different trajectories, as reflected by the connectivity maps they generate at different time points. Figure 6D quantifies the models’ performances at all measured time points (a subset of which is shown in panel A), by showing the density of synaptic connections during development. It shows that the single epoch model is far from the empirical values at multiple time points, whereas the accuracy of the multi-epoch one is consistent throughout worm development (see Fig. S7 for the average over data splits). Interestingly, the values of the probabilities that the multi-epoch model assigns to individual synapses are higher than those of the single epoch model at early stages. Fig. 6E, which shows the likelihood ratio of the models over the different connectomes across development, demonstrates that the multi-epochs model is consistently better by a large margin in the early developmental stages. Thus, multiplicity of developmental epochs emerges as a critical component for building the worm connectome.

**Fig. 6.**
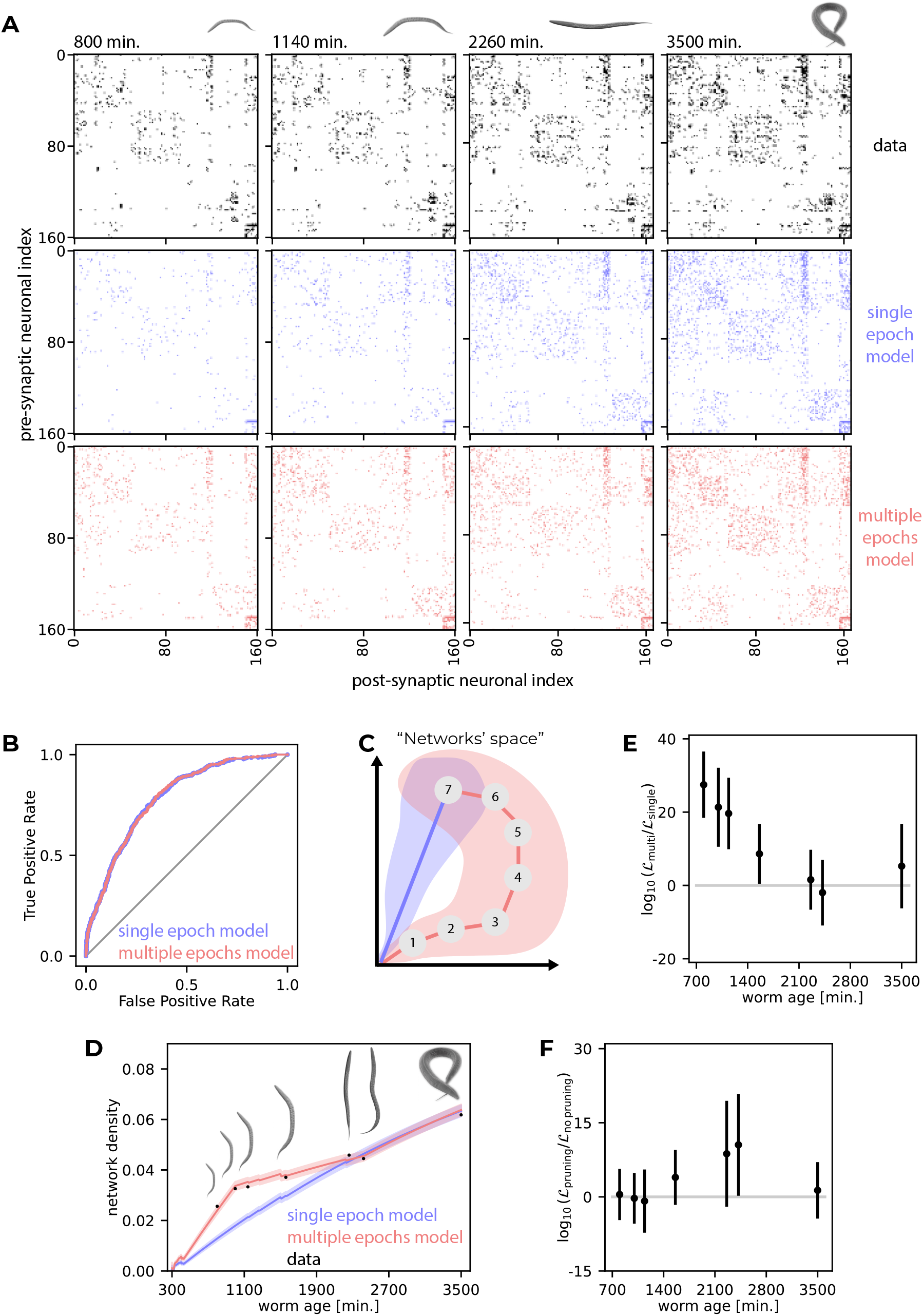
Multiple developmental epochs and synaptic pruning are necessary to follow the observed developmental path. **A**. Connectivity maps of data (top), single draws of the single epoch model (middle), and the multi-epoch model (bottom) at time points corresponding to worms 1, 3, 5 and 8 (adult) in [16]. **B**. ROCs of the single and multi-epoch models at adult age on test data. **C**. An illustration of a possible scenario where the single and multi-epoch models follow disparate paths in the space of networks but end up in the same place. **D**. Network density versus worm age for the measured connectivity maps (test data), single epoch model, and multi-epoch model. Solid lines represent the models’ average density, and shaded areas the 1-std regions. Calculations are exact (see Methods). **E**. Average log-likelihood-ratios of the multi-epoch and single epoch models over 20 splits of the data into training and test subsets. Error-bars show 1-std. **F**. Average log-likelihood-ratios of the multi-epoch model with synaptic pruning and the multi-epoch model with no pruning over 20 splits of the data into training and test subsets. Error-bars show 1-std.

To explore the role of synaptic pruning in connectome development, we also studied the performance of a model that uses multiple epochs without pruning. We found that pruning seems to improve the models’ ability to reproduce the synaptic connectivity maps at different developmental stages (Fig. 6F).

We note that since the reconstructions at different time points are not of the same animal, it is not clear whether there is a biologically-realistic (rather than a mathematical) developmental “path” that would go through them. The fact that a model based on three epochs is sufficient to create such a path suggests that individual variability does not imply divergent wiring diagrams, and that connectomic variability may be curbed by the developmental plan.

### The generative model reveals the stable “backbone” of the connectivity map

The scarcity of connectomic data makes it difficult to study individual diversity in circuits’ structure and function. Importantly, our generative models for the connectomes are probabilistic by nature, and so, given their accuracy, they can be used to explore the variability and shared connectomic structures, and can be compared to the handful of examples where the same circuits have been reconstructed in different animals [16, 45–47]. We quantified the variance of the connectomes generated by the generative models by the expected normalized Hamming distance between model runs, namely, the average fraction of synapses that are formed in one run but not in the other (see Methods). We compared this value to the cross-variance of the model and the data, similarly defined as the expected normalized Hamming distance between a run of the model and the data. We find that the values of cross-variance of the model and the data during the different stages of development lie within 1 std of the model’s variance from its mean (Fig. 7A, Fig. S8A). Thus, the empirical data is consistent with typical model-generated connectomes.

**Fig. 7.**
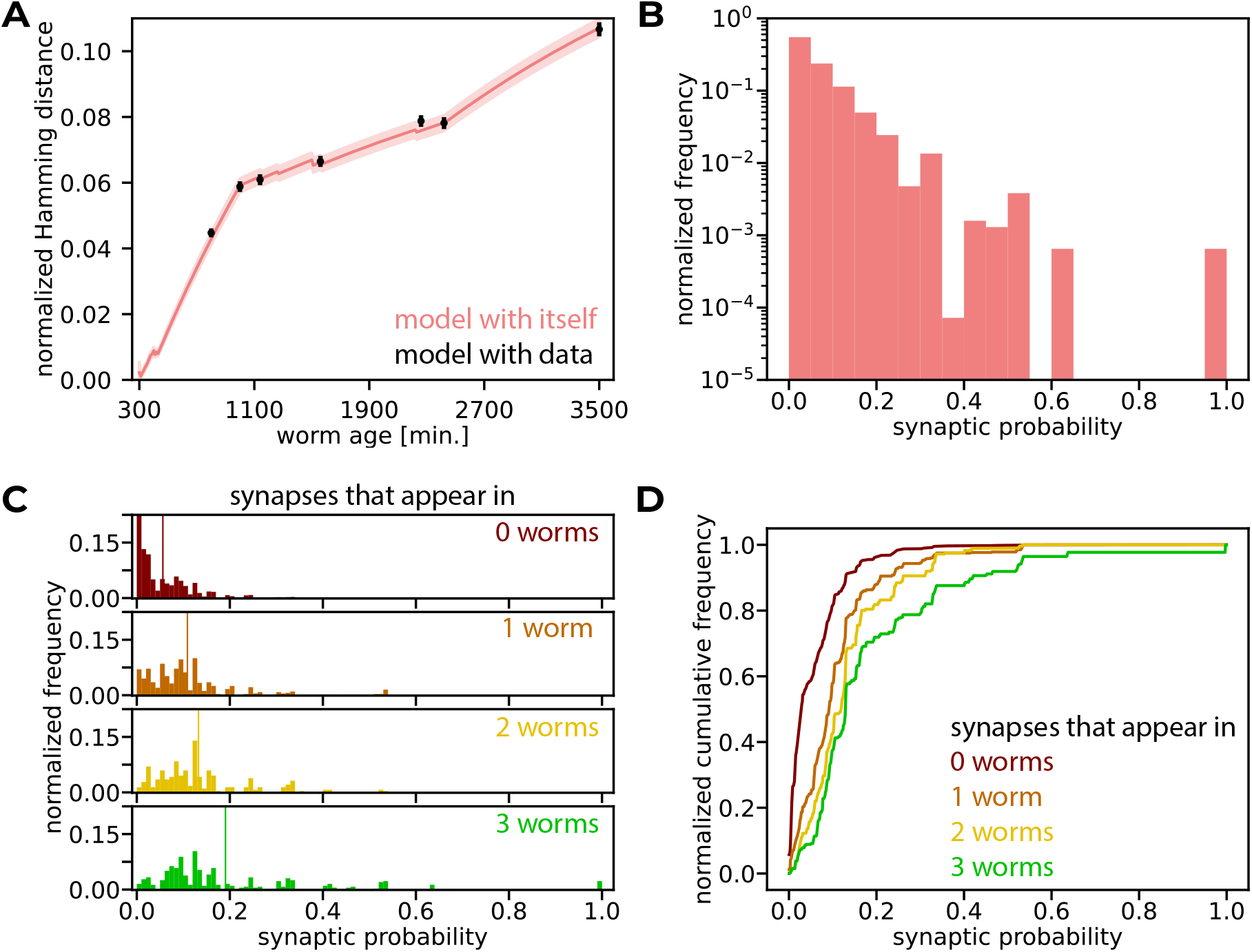
The model reveals the stable backbone of the connectivity map. **A**. The variance of the model (coral line) and the cross-variance of the model and the data (black scattered dots), measured by normalized Hamming distance. The solid line represents the mean normalized Hamming distance between model runs, and the shaded area is the 1-std region. The dots represent the mean normalized Hamming distance between a run of the model and the data, and the error-bars denote 1-std. Calculations are exact (see Methods). **B**. The normalized histogram of the probabilities the model assigns to all possible synapses. **C**. Normalized histograms of synaptic probabilities predicted by the model, when grouped by the number of datasets in which they appear. Vertical lines show the distributions’ means. **D**. Normalized cumulative distributions of synaptic probabilities predicted by the model, when grouped by the number of datasets in which they appear.

Ideally, we would next compare the variance of the model to the biological variance of connectomes measured experimentally. However, the limited number of reconstructed connectomes means we can only compare the two adult worms reconstructed in [16], which show a normalized Hamming distance of 0.04, compared to 0.105 *±* 0.002 (mean *±* std over 20 splits) of the model’s. Additional reconstructed connectomes would eventually allow the quantification of the natural variance and the divergence of the model’s variance from it. As a proxy for estimating the biological variance, we group the synapses according to the number of experimental data sets in which they appear. We here use, in addition to the two adult worms reconstructed in [16], also the reconstructed connectome from [45–47]. The distribution of the probabilities the model assigns to synapses spans the whole range of [0, 1], with most of the synapses assigned very low probabilities (Fig. 7B, Fig. S8B). We then compared the model’s probabilities to the prevalence of synapses in the available data. Figure 7C shows the normalized frequencies for 4 sets of synapses: those that appear in all 3 connectomes, those that appear in 2 out of 3, etc. Figure 7D then shows the corresponding cumulative frequencies and the different nature of these classes of synapses (see also Fig. S8 C,D for the average histograms across data splits). Although the generative model was trained on just one of the connectomes, synapses that exist in more data sets are distinctly predicted to exist with higher probability. Moreover, despite the fact that only *∼* 6% of the possible synapses exist in the connectome we trained on, the model predicts that some synapses will be there with a probability close to 1, and these indeed appear in all the 3 data sets we used to test the model. Once more data sets become available, future models will be able to identify more “certain” synapses that appear in all worms and allow us to characterize the variability of others.

## Discussion

We presented a family of generative models for the development of connectomes and used them to study *C. elegans*. These models suggest that a surprisingly small number of biological and physical features may govern the construction of the nervous system of the worm, as our compact models accurately predicted individual synaptic connections, neuronal degree profiles, and sub-network motifs during development. These developmental models of the connectome allowed us to show that while it is possible to construct an accurate connectome of an adult nematode with a single set of developmental parameters, this would result in the creation and pruning of significantly different numbers of synapses during development than the observed ones. Our results suggest that multiple developmental stages and synaptic pruning are necessary for following the empirical measurements of development. Importantly, our models also show that construction rules that rely on the common classification of neurons into types are highly redundant. Indeed, we identified a significantly smaller number of neuronal cell types and simple construction rules for these inferred cell types, which yield more accurate connectomes.

The models are immediately related to the classic ideas of molecular “lock and key” mechanisms [26], morphological rules, etc. Thus, our models provide a means to synthesize connectomes throughout development, and indicate a possible algorithmic way for biology to implement them. Yet, while our models suggest biologically plausible feature-based construction rules for building the lion’s share of a full nervous system, the real wiring rules employed in the worm may be different. However, we emphasize that the accuracy of our models imply that even if that is the case, the rules of our models and the real ones highly overlap in terms of their predictions. Thus, our model offers a draft of the construction rules that would need to be verified experimentally, or different features that are incorporated to implement equivalent rules would need to be identified.

The small number of features that our models rely on and the stochastic nature of our model mean that connectomes that are synthesized by it (i.e., multiple “draws” from the model) would differ from one another. While adding construction rules could allow for lower variability of the resulting connectomes, this would probably require genetic encoding of these rules as well as the biophysical mechanisms that would be needed to implement them accurately and to correct errors. We therefore submit that the simplicity of our model and its small number of features is particularly appealing, as the variability that our models predict is consistent with their characteristic variability from the data. Moreover, our models assign higher probabilities to synaptic connections that are shared across individuals. Thus, our models imply a connectomic “backbone”, which is highly conserved across individuals, and volatile parts. With the reconstruction of more wiring diagrams of individuals, and potentially characterizing the detailed neural activity of these reconstructed circuits, our framework could be extended to explore the implications of the different construction rules on function, as well as individual variability of neural networks and of the resultant behavior.

It is important to note that the data used here to learn the models were obtained from different individuals, and so, it is not clear that the “developmental path” we assumed by considering the different worms as time points along one trajectory would be realizable. However, the fact that we were able to train models to follow a developmental path across individuals is a testament to the strength of the models and implies a relatively high robustness of developmental trajectories. As reconstructions of connectomes of single animals across development are beyond current experimental ability, testing these ideas would require new experimental and computational methods.

Other natural extensions of our work are theoretical and experimental explorations of whether the dynamic developmental parameters arise from changes in demands and resources during the growth of *C. elegans* or from functional requirements of the developing circuit. Moreover, as synaptic pruning can be thought of as an error-correction mechanism, reminiscent of kinetic proof-reading [48], it would be interesting to characterize its specificity as well as to investigate whether it can be viewed as balancing energetic efficiency in constructing the circuit and the cost of the regulation needed to build the correct connectivity map.

Finally, beyond *C. elegans*, our results suggest a general framework for studying the construction of connectomes and their underlying design principles. Such computational models, and their interpretable features, as we suggest here, would lay the infrastructure for understanding the design of neural connectivity, which would be instrumental for interacting with biological neural networks, building similar ones, re-designing and fixing them.

## Methods

### Data pre-processing

The connectomes are based on [16], resulting in connectivity maps between 180 neurons of the nerve ring in 8 nematodes at 7 developmental stages (2 of the nematodes are adults). The analysis focuses on chemical synapses (and not electrical gap junctions), and ignores auto-synapses (synaptic connections from a neuron to itself). Synaptic connections are binarized (a connection either exists, i.e, there is at least one synapse in the data between the corresponding neurons, or it does not exist). Neuronal positions are obtained from [39] and birth times from [38]. To resolve a discrepancy between the birth times of neurons and the subsets of neurons in the different connectomes, when training and analyzing models across development, 13 neurons are omitted. When training in a single developmental epoch, the eighth data set is considered as the test set and is not used for training. When training in multiple developmental epochs, all the possible synapses of the adult worm are split randomly into two halves, one of which is used for training and the other for testing (data sets 2, 6 and 8 are used for training to define 3 developmental epochs). As the training procedure of the model that includes reciprocity (see model specification) requires that both synapses of each dyad appear in the data set, half the dyads are randomly sampled for the training data (namely, half the possible unordered sets of pairs of neurons), and both of the synapses of each dyad are included. Models are trained using 20 splits of the possible synapses into training and test subsets. The ordering of neurons in adjacency matrices is according to their types, and within a type the ordering is alphabetical.

### A developmental model for the connectome of *C. elegans*

The model grows a network representing the nervous system of *C. elegans* in discrete time steps of 10 minutes each. At every time step, 3 operations are performed to change the architecture:

1. Neurons are born if their birth time [38] has passed since the last step (in the past 10 minutes). Each neuron is given coordinates according to its 3D location in the worm’s body [39]. The coordinates are normalized to the length of the worm (800*µm*, [39]). Additionally, each neuron is given a neuronal cell type.
2. Synapses are added with a probability that depends on their types and the distance between their neurons. The type is induced by the types of the pre-synaptic and post-synaptic neurons: (*τ*_*pre*_, *τ*_*post*_) and the distance is the normalized Euclidean distance between them: 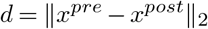, where *x* is a 3-dimensional coordinate vector of a neuron. Each synaptic type has a specific baseline formation probability 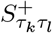 (where *k* and *l* stand for the indices of the pre- and post-synaptic neuronal cell types, respectively), which decays exponentially with the distance between neurons according to a decay rate *β*_*t*_. To model the elongation of the worm during its development, *β*_*t*_ is increased with time, while neuronal locations are kept constant. The growth of *β*_*t*_ is piecewise linear and results in a 16-fold elongation in total. Linear growth periods connect developmental stages with measured average lengths [49]. In this manner, an isotropic growth is modeled [16]. All in all, the probability for synaptic formation at time *t* is given by 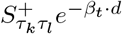 Every non-existing synapse is formed with the corresponding probability independently of other synapses.

Existing synapses are pruned independently of one another with constant probability *S*^*−*^.

### Calculation of the likelihood of a network

As synapses are formed and pruned independently of one another, the probability that an output of the model, *G*, with parameters 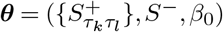, equals the data *G*^*data*^ is given by:

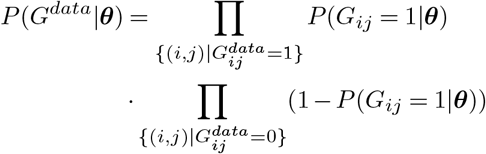

The probability of a synapse in the network to exist at time *T* since the birth of the youngest of the neurons that form it can be calculated recursively by the relation:

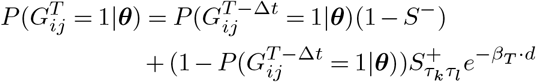

and the initial condition,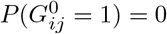 where *G*^*T*^ is the adjacency matrix of the network at time *T*, Δ*t* is the time step (10 min.), and the rest of the parameters are as described in the model specification.

### Model training procedure (single developmental epoch)

The model is controlled by a 3-tuple of parameters,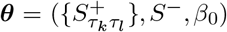 To fit them, a grid search is performed over 1600 (40 *×* 40) values of *S*^*−*^ and *β*_0_, considering a range of [0, 0.05) for *S*^*−*^ and *β*_0_. For each pair 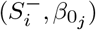, a set of binary searches is performed, one for each synaptic type (*τ*_*k*_, *τ*_*l*_) (defined as a pair of neuronal cell types), separately from one another. The objective of the search is to find the value of 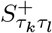 that will result in an average number of synapses of type (*τ*_*k*_, *τ*_*l*_) that matches the data (up to a tolerance of 0.05). The average number of synapses of type (*τ*_*k*_, *τ*_*l*_) is given by:

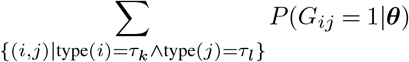

where *G* is the output of the model. The probabilities in the sum are calculated as described in the calculation of the like-lihood. As the number of synapses is monotonic in 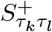 (given a pair of values for *S*^*−*^ and *β*_0_), the binary search is guaranteed to keep approaching the objective. If the objective is not achievable, the search terminates once the value of 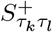 gets close enough to its bounds (0 and 1). The termination condition used here is a difference of less than 10*e−* 10. The procedure above results in 1600 3-tuples of model parameters that conserve the number of synapses of each type in the data (if possible). For each set of parameters, the log-likelihood of the data is computed (as described above), and the set with the maximal value is chosen.

### Models based on subsets of features

#### Birth-time model

In this model, as described above, all neurons have the same cell type (a single *S*^+^ value), and there is no dependence on distances between neurons (i.e., setting *β*_0_ = 0).

#### Distance model

In this model, the distance decay matrix, defined as

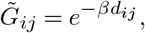

where *d*_*ij*_ is the distance between neurons *i* and *j*, is normalized with a global normalization factor *λ* such that

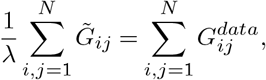

where *G*^*data*^ is the adjacency matrix of the data and *N* is the number of neurons (i.e., conserving the density of the data). To fit the model parameter *β*, multiple values in the range of [0, 4) are considered, and the one yielding the maximum likelihood of the data is chosen.

### Greedy clustering of neurons into types

Neuronal cell types are aggregated in a greedy manner. The objective function of the greedy clustering is referred to as “hit rate”, and is defined as the expected synaptic agreement with the data when sampling uniformly the “correct” number of synapses of each type (i.e., the same number of synapses as in the data). The optimization process intends to maximize the hit rate. Initially, each neuron is given a unique type. Then, iteratively, all merging possibilities of a pair of types are examined, and the one resulting in the smallest decrease in the hit rate is performed, reducing the number of neuronal cell types by 1. An explanation of the calculation of the hit rate follows.

It holds that

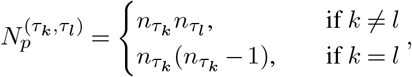

where 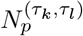 is the number of possible synapses of type (*τ*_*k*_, *τ*_*l*_) and 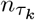 is the number of neurons of type *τ*_*k*_. The difference between the cases is due to the omission of auto-synapses.

The number of existing synapses of type (*τ*_*k*_, *τ*_*l*_) in the data is denoted by 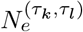, and, as described above, this number of synapses is assumed to be sampled uniformly. Thus, the hit rate for this synaptic type is given by

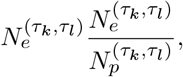

as the probability of each sampled synapse to appear in the data as well is

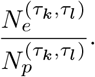

All in all, the total hit rate is given by

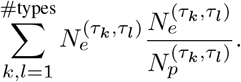

### Validation of not over-fitting using 8 inferred neuronal cell types

To verify that the choice of 8 types does not result in over-fitting, a single draw of a trained model with a known ground truth of 8 neuronal cell types (a “synthetic connectome” with 8 types) was sampled and used as a control data set to train a new, second, model. This second model was trained using a number of neuronal cell types ranging from 1 to 15, where it is expected to see an improvement in performance on test data when using up to 8 types, and a decrease thereafter (as a result of over-fitting). 100 different “synthetic connectomes” were used as test data, and this classical over-fitting pattern emerged. However, the model that was trained on the experimentally measured data using 8 types keeps improving as the number of types increases (i.e., even above 8), suggesting that the use of 8 types does not result in data over-fitting (see Fig. S2B).

### Compact model representations

Compact representations of models were found using quantization. Only the 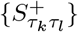 values were quantized (where values of 0 were forced to remain 0 after quantization). Formally, a quantization into *K* values is a function 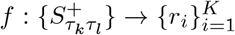. Such a quantization would result in a compact representation of the model, with *K* + 2 parameters: *K* values for the quan-tized 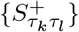 values, *S*^*−*^ and *β*_0_. The compact model is then identical to the full one, except for using 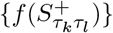as base-line synaptic formation probabilities.

The sum of squared errors of the quantization is used as a measure for dissimilarity of the compact set of values and the original set of values. It is defined as

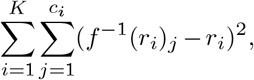

where 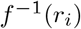 is the set of *S*^+^ values mapped to *r*_*i*_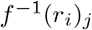 is the j-th element in this set and *c*_*i*_ is the cardinality of this set.

Given a partition of 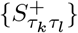 into *K* subsets 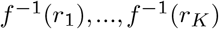, the values 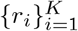 that would minimize the sum of squared errors are

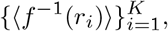

and the sum of squared errors becomes

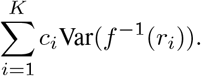

Thus, the problem of finding the best quantization into *K* values is equivalent to the problem of finding the partition of 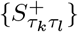 into *K* subsets such that the weighted sum of their variances is minimal.

This problem can be solved using Dynamic Programming. Given a set of *n* numbers, 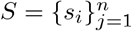, algorithm 1 finds their split into *K* non-empty subsets with the minimal sum of weighted variances, in polynomial time. It uses the following notations:

- The function **Sort** gets a set *S* of *n* numbers and returns them as a sorted *n*-tuple (*x*_1_, …*x*_*n*_). Namely,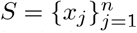, (set equality), and ∀ 1 *≤ j < j*^*′*^ *≤ n*, it holds that *x*_*j*_ *≤ x*_*j*_*′*.
- *V* is a lookup table of which keys are tuples (*i, j*), where 1 *≤ i ≤ K*, 1 *≤ j ≤ n, i ≤ j*, and values are tuples (*v, P*), where *v* is the minimal weighted sum of variances of a partition of the *j* first numbers into *i* non-empty subsets, and *P* is the partition resulting in the minimal variance (a set of sets).
- {{*x*}} is a set containing the set {*x*}, and similarly {{*x*},{*y*}} is a set containing the sets {*x*} and {*y*}.

Binary search is used to find an accurate compact representation of a model. Accuracy is measured by the fraction of the AUROC value of the compact model from the full one. The objective of the search is to find the maximal unexplained variance fraction for which the accuracy condition is satisfied. The search halts when both the performance condition is satisfied and the size of the interval of search is below a threshold (a threshold of 10*e−* 10 is used).

### Graph structural feature calculations

For calculations of graph structural features, the following formalism is used: each synapse can be thought of as a random variable, denoted *X*_*ij*_, distributed Bernoulli with a probability *p*_*ij*_ for success, where [*p*]_*ij*_ is the average matrix of the model. A single sample of the model is composed of the values of *X*_*ij*_ for all 1 *≤ i < j ≤ N*, where *N* is the number of neurons.

- Network density: the number of synapses in the network is given by

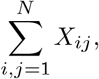 so the network density is

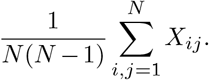

#### Algorithm 1

Find the partition with the minimal weighted sum of variances

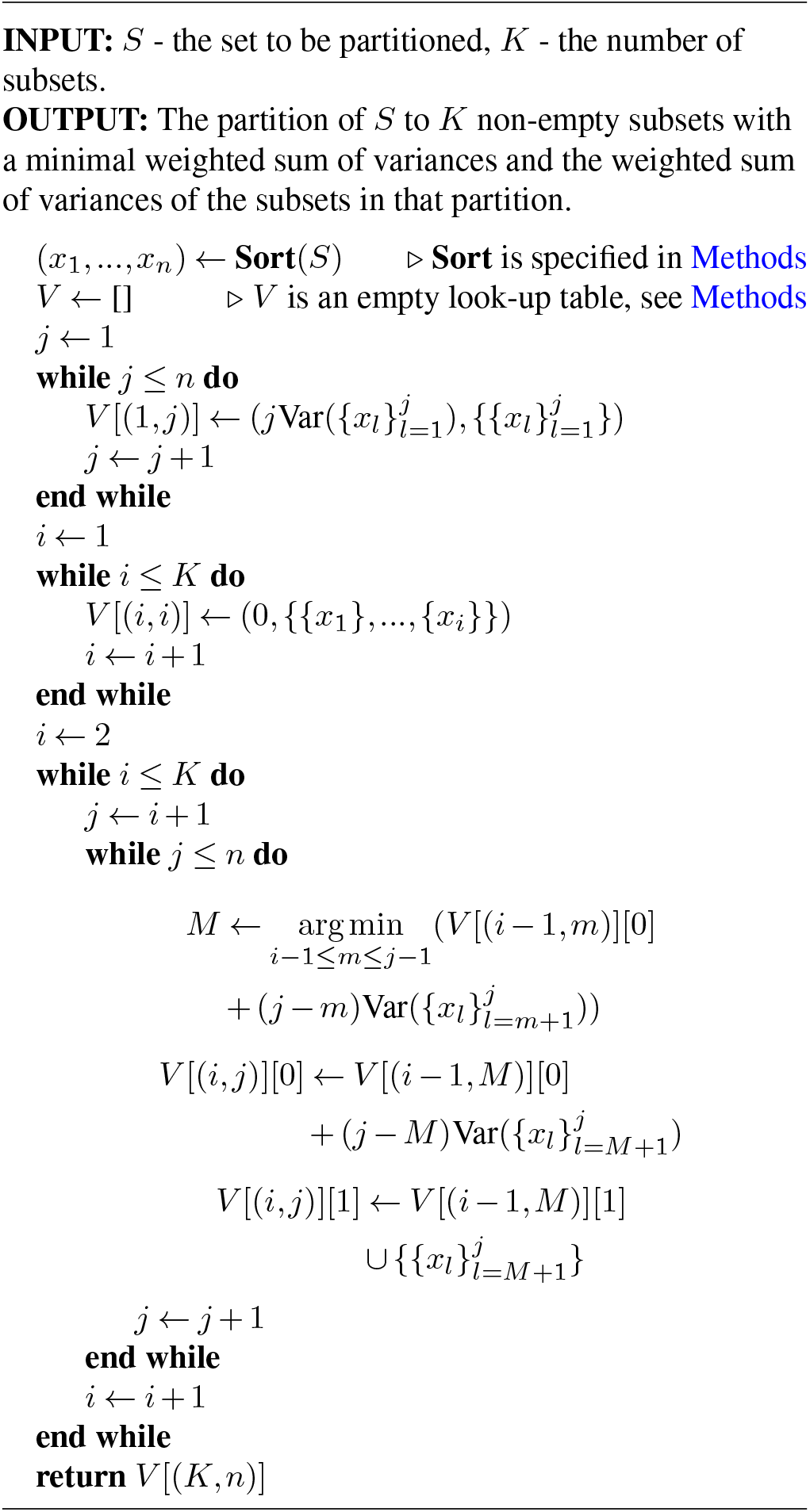

Thus, the average network density is given by

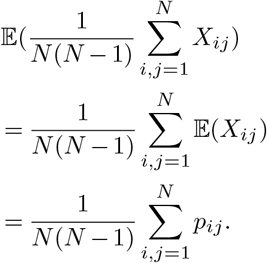

The variance is similarly

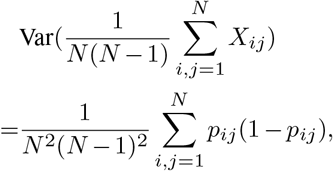

where the equality holds when 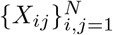 are independent. The standard deviation is then

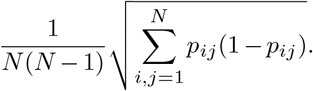

For the reciprocity dependent model, the covariance of the dyads should be added to the variance, which yields

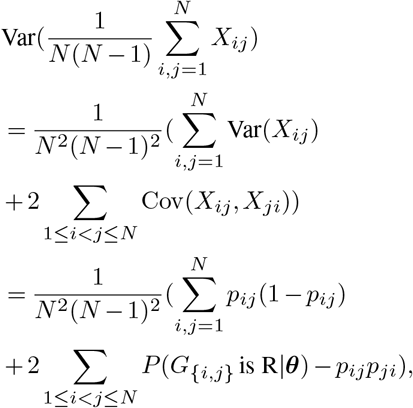

where *G* is the output of the model, {*i, j*} is an unordered set of neurons and *R* stands for Reciprocal (namely, both synapses of the dyad are formed).

- Degrees: in and out-degrees are calculated similarly, using the transposed average matrix of the model. Thus, only the calculations for in-degrees will be explained. The in-degree of the j-th node of a graph is given by

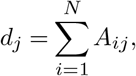 where *N* is the number of nodes in the graph and [*A*]_*ij*_ is its adjacency matrix. Thus, in the case discussed here, the in-degree of the j-th neuron in the model can be defined as a random variable that takes values between 0 and *N* according to

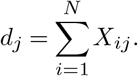

Such a probability distribution (which is a result of a summation of *N* independent Bernoulli random variables with success rates of 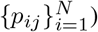 is called Poisson-Binomial and its probability mass function can be calculated using the characteristic function [50]. Note that this holds for the reciprocity dependent model as well, as only reciprocal synapses are dependent and they cannot share neither a row nor a column. The expected value and variance are given by

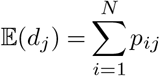 and

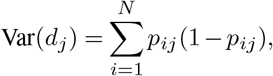 respectively.
- Histogram calculations: given *m* bins with edges 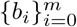 and *l* independent random variables 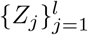 that take values in [*b*_0_, *b*_*m*_), the indicator 𝟙_*ij*_ is defined as

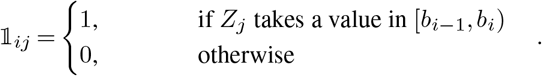

The frequency of *Z* values in the *i*-th bin is then

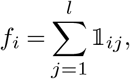 which distributes Poisson-Binomial, and its expected value and variance are

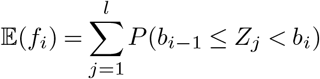 and

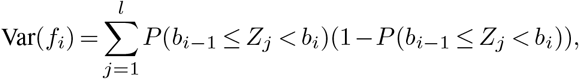 respectively. The cumulative histogram is defined as

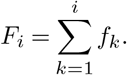

Its expected value is given by

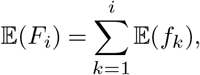 and is variance can be calculated as follows:

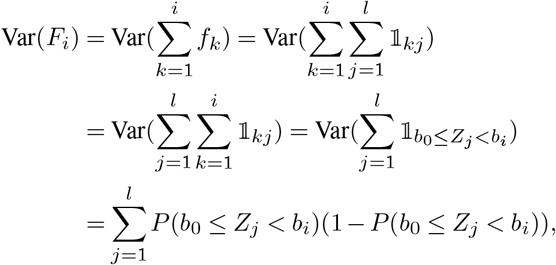 where the last equality holds because 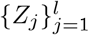 are independent. For degree distribution histograms, bin edges were taken as

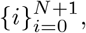 and the random variables are the degrees of the neurons:

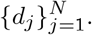

For the histogram of distances between connected neurons, the distances at adulthood were considered. Bin edges were taken as

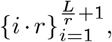 where *r* is the resolution (0.1[*µm*], which is less than the minimal possible distance between neurons, which is *∼* 0.7[*µm*]), and *L* is the maximal distance between neurons, which is *∼* 776[*µm*]. The random variables were defined as

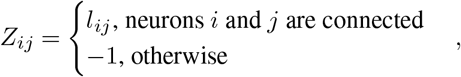 where *l*_*ij*_ is the distance between neurons *i* and *j*. Similarly, for birth time histograms, bin edges were taken as

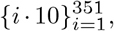 where the resolution of neuronal birth times in the data is 10[min .], and the age of an adult nematode is 3500[min .]. The random variables were defined as

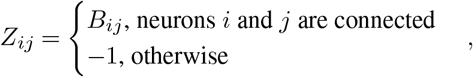 where *B*_*ij*_ is the maximal birth time of the neurons *I* and *j*.
- Triad distribution: the triad distribution of a graph is calculated by simply counting the number of triads of each one of the 16 types, and normalizing all of them by the total number of triads in the graph (which is 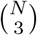). The model’s average triad distribution and its standard deviation were calculated numerically: 1000 model samples were generated and the distribution for each one of them was calculated using the networkx package for Python, version 3.2.1, resulting in 1000 vectors **v**^*k*^ of length 16. The *i*-th entry of the mean distribution is then

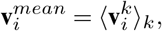 and the *i*-th entry of the standard deviation vector is

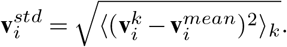
- Network reciprocity: the number of possible reciprocal dyads in the network is given by 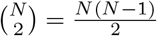. Let *Y*_*ij*_ be a new random variable that takes 1 when a dyad is reciprocal and 0 otherwise (and thus distributes Bernoulli), defined as *Y*_*ij*_ = *X*_*ij*_*X*_*ji*_. When synapses are independent, it holds that the success rate is *p*_*ij*_*p*_*ji*_. When reciprocal synapses are dependent, the success rate becomes *P* (*G*_{*i,j*}_ is R|***θ***), where *G* is the output of the model, {*i, j*} is an unordered set of neurons and R stands for Reciprocal (namely, both synapses of the dyad are formed). It can be calculated as described below. The total number of reciprocal dyads is then

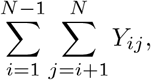 and the reciprocity is thus (after a proper normalization)

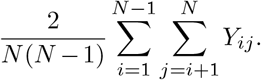 The average reciprocity is then

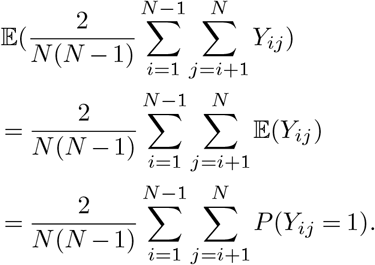

The variance is similarly given by

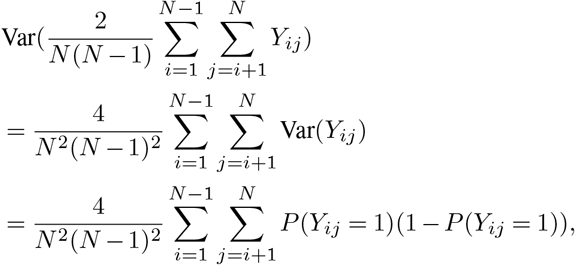 so the standard deviation is

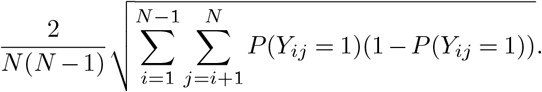

### Reciprocity-dependent models

The model that incorporates statistical dependence of reciprocal synapses is identical to the model specified above, except for the second operator. Here, synapses are added with a probability that depends on the state of their reciprocal synapse as well. The probability becomes min 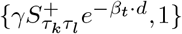 when the reciprocal synapse exists, where *γ* is an additional parameter of the model that controls the tendency of the model to increase its reciprocity. The likelihood of the model is now calculated by iterating the states of all dyads. A dyad has 4 possible states: reciprocal - both synapses are formed, upper - only the synapse of the upper triangle of the adjacency matrix is formed, lower - only the synapse of the lower triangle of the adjacency matrix is formed, and empty - no synapse is formed. The states are denoted in short R, U, L and E, respectively. The likelihood is then given by:

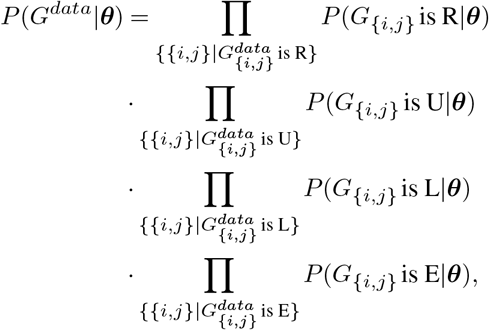

where *G* is the model adjacency matrix, *G*^*data*^ is the data adjacency matrix and {*i, j*} is an unordered set of indices (a dyad).

The probability predicted by the model of a dyad being in a specific state can be calculated recursively. Following is the recursive relation for the reciprocal state as an example, and the rest of the states are calculated similarly.

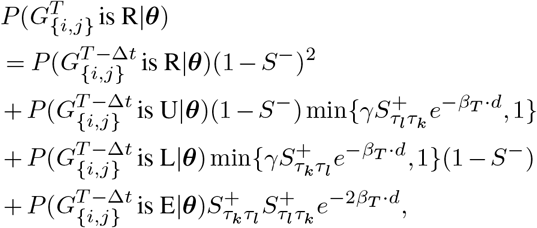

where 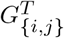 is the state of the dyad {*i, j*} at time *T, d* is the distance between the neurons, *τ*_*k*_ is the type of neuron *i, τ*_*l*_ is the type of neuron *j*, and *i < j* is assumed. The initial condition for the recursion is

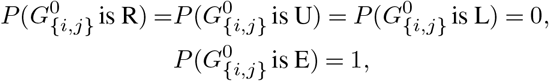

as all dyads are empty at first.

The training of models is similar to a single epoch model training procedure, but here the *S*^+^ values for the 2 synaptic types of a dyad are found simultaneously. There are 2 binary searches that are alternately updated. After each update, the expected number of synapses of both types are compared to the numbers of synapses of these types in the data. The value of *γ* is set constant during training, and different models are trained for different values. If the simultaneous search does not converge, the search halts when the window size of both searches is smaller than 10^−10^.

### Model training with multiple developmental epochs

The training procedure here is similar to the one described in the single epoch model training procedure section. Multiple epochs models allow 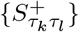 values to change between discrete developmental stages. Thus, for each synaptic type (*τ*_*k*_, *τ*_*l*_), several subsequent binary searches are necessary to determine the values for 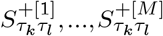, where *M* is the number of epochs. The *m*-th value is found using a binary search whose objective is to match the number of synapses in the data of the nematode at developmental stage *m*, on average. The previously found values for 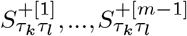 are used to calculate the probability of each dyad to be at a specific state using recursive relations. Following the example above, the probabilty of a dyad to be reciprocal becomes

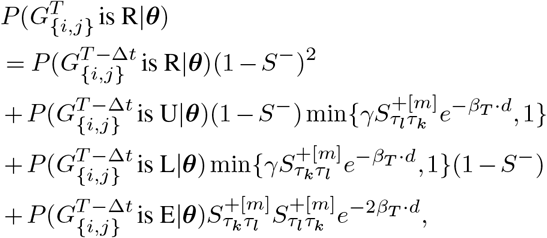

where 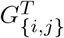 is the state of the dyad {*i, j*} at time *T, m* is the developmental stage at time *T, d* is the distance between the neurons, *τ*_*k*_ is the type of neuron *i, τ*_*l*_ is the type of neuron *j*, and *i < j* is assumed. The initial condition for the recursion is

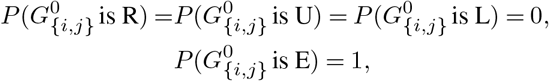

as all dyads are empty at first.

### Model variance calculation

The variance of the model is defined as the expected normalized Hamming distance between two model runs, namely, the fraction of flipped synapses. The expected number of flipped synapses of an individual dyad {*i, j*} (an unordered pair of neurons) can be defined as a random variable, denoted *X*_{*i,j*}_, that takes values from {0, 1, 2}. As discussed above, each dyad has 4 possible states (R, U, L and E). Thus, there are 4^2^ = 16 possible states for a dyad in 2 different runs. An enumeration over the 16 possible states yields the probabilities for *X*_{*i,j*}_ to take each one of its possible values (e.g., *X*_{*i,j*}_ = 2 for the states (R, E), (E, R), (U, L), (L,U)). The probability of a dyad to be in a certain state is calculated as described above. The variance of the model is then given by

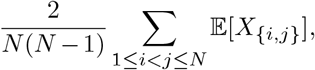

and its standard deviation by

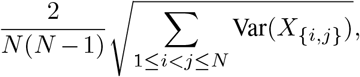

as the dyads are statistically independent.

The cross-variance of the model and the data is defined similarly as the expected normalized Hamming distance between the model and the data. The calculations and definitions here are identical, but the distribution of the pair of dyadic states over the 16 possibilities is different (as the state of the dyad in the data is deterministic).

### Neuronal birth time noising

The birth time of each neuron, *b*_*i*_, was added with noise *ϵ*_*i*_, sampled uniformly from [*−δw*_*i*_, *δw*_*i*_), where 0 *≤ δ ≤* 1 is the noise level and

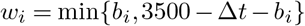

is the sampling window size (3500 minutes is the age of a adult worm), and Δ*t* is the time step of the model (which is 10[*min*.]). Thus, the noised birth time, 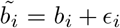, is guaranteed to not exceed the interval [0, 3500 *−* Δ*t*), namely, the worm’s developmental time window (the upper bound of the interval is 3500 *−* Δ*t* and not simply 3500 to allow all neurons to form connections during at least one step), and the expected value of the noise is 0.

10 noise levels from 0.1 to 1 were considered (with a difference of 0.1 between subsequent levels), and for each level birth times were noised 100 times.

## ACKNOWLEDGEMENTS

We thank Adam Haber, Gal Goldman, the rest of the Schneidman lab members and Meital Oren-Suissa for discussions, comments and ideas. This work was supported by Simons Collaboration on the Global Brain grant 542997 (ES), Israel Science Foundation grant 137628 (ES), Israeli Council for Higher Education/Weizmann Data Science Research Center (ES), Martin Kushner Schnur, and Mr. & Mrs. Lawrence Feis. ES is the incumbent of the Joseph and Bessie Feinberg Chair. This research was also supported in part by grants NSF PHY-1748958 and PHY-2309135 and the Gordon and Betty Moore Foundation Grant No. 2919.02 to the Kavli Institute for Theoretical Physics (KITP).

## Supplementary figures

**Figure S 1.**
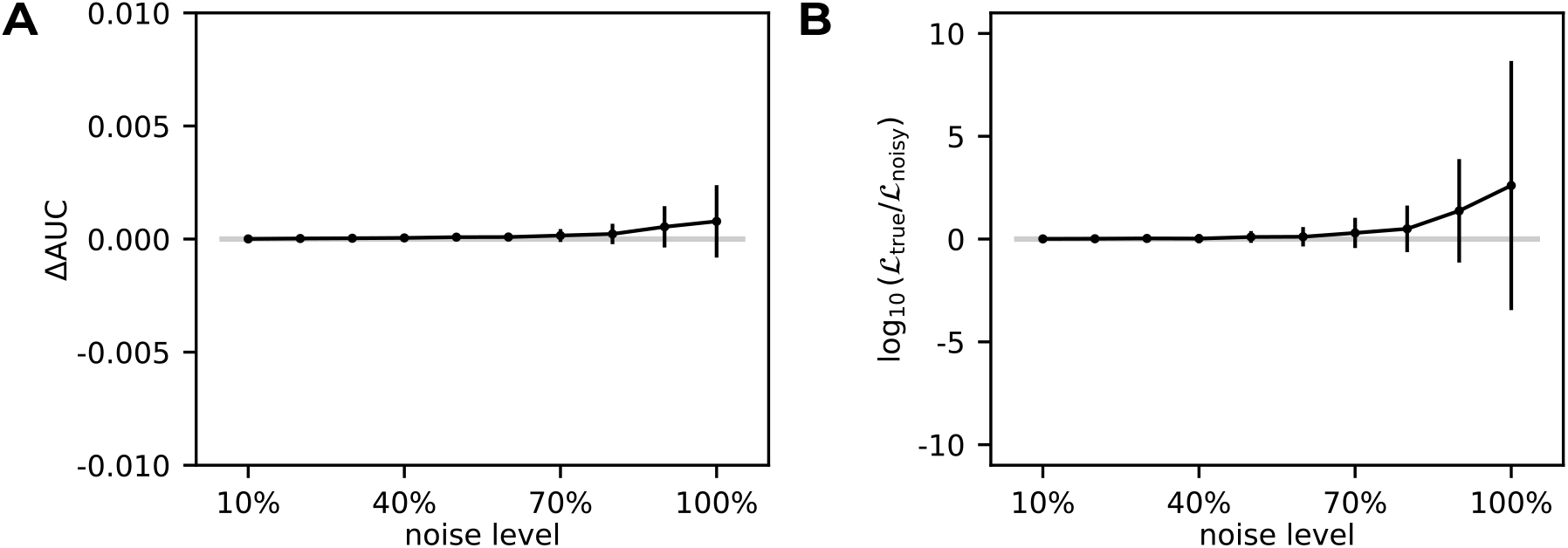
**A**. The average difference between the AUROC on test data of a model that uses true birth times of neurons and that of models that use a noised version of them versus the level of noise (see Methods). Averages are over 100 different additions of noise, and error-bars show std. **B**. The average log likelihood ratio on test data between a model with true birth times and models with noised birth times versus the level of noise. Averages are over 100 different additions of noise, and error-bars show std. Calculations are exact (see Methods).

**Figure S 2.**
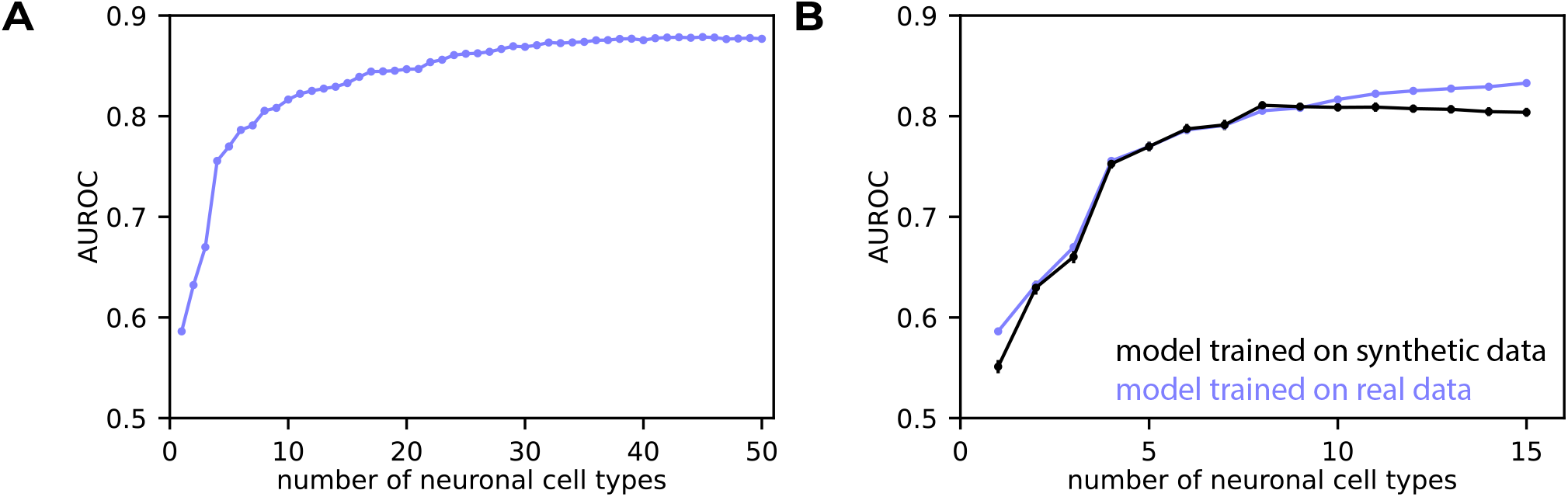
8 inferred neuronal cell types do not result in data over-fitting. **A**. Model AUROC versus the number of inferred cell types used. **B**. Model AUROC versus the number of inferred cell types used. Purple - a model that was trained and tested on data (like in **A**). Black - a model that was trained and tested on outputs of a pre-trained model that used 8 inferred cell types (see Methods). Error-bars show standard deviation over different model samples serving as test data, and are smaller than the marker size for most dots.

**Figure S 3.**
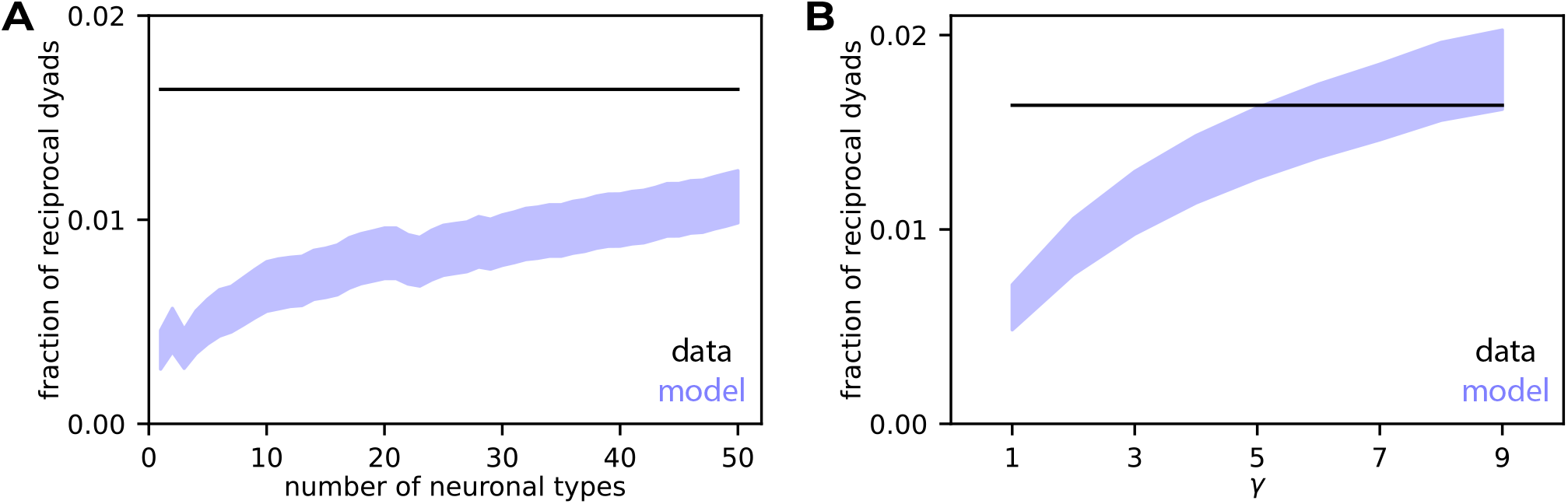
A simple mechanism for controlling reciprocity is sufficient to reproduce the observed value. **A**. Reciprocity fraction versus number of neuronal cell types when synapses are formed independently of one another. The shaded area represents the 2-std region of models. Calculations are exact, see Methods **B**. Reciprocity versus *γ*, the factor increasing the probability to form a synapse given its reciprocal synapse is formed, for a model with 8 types. The shaded area represents the 2-std region of models. Calculations are exact, see Methods.

**Figure S 4.**
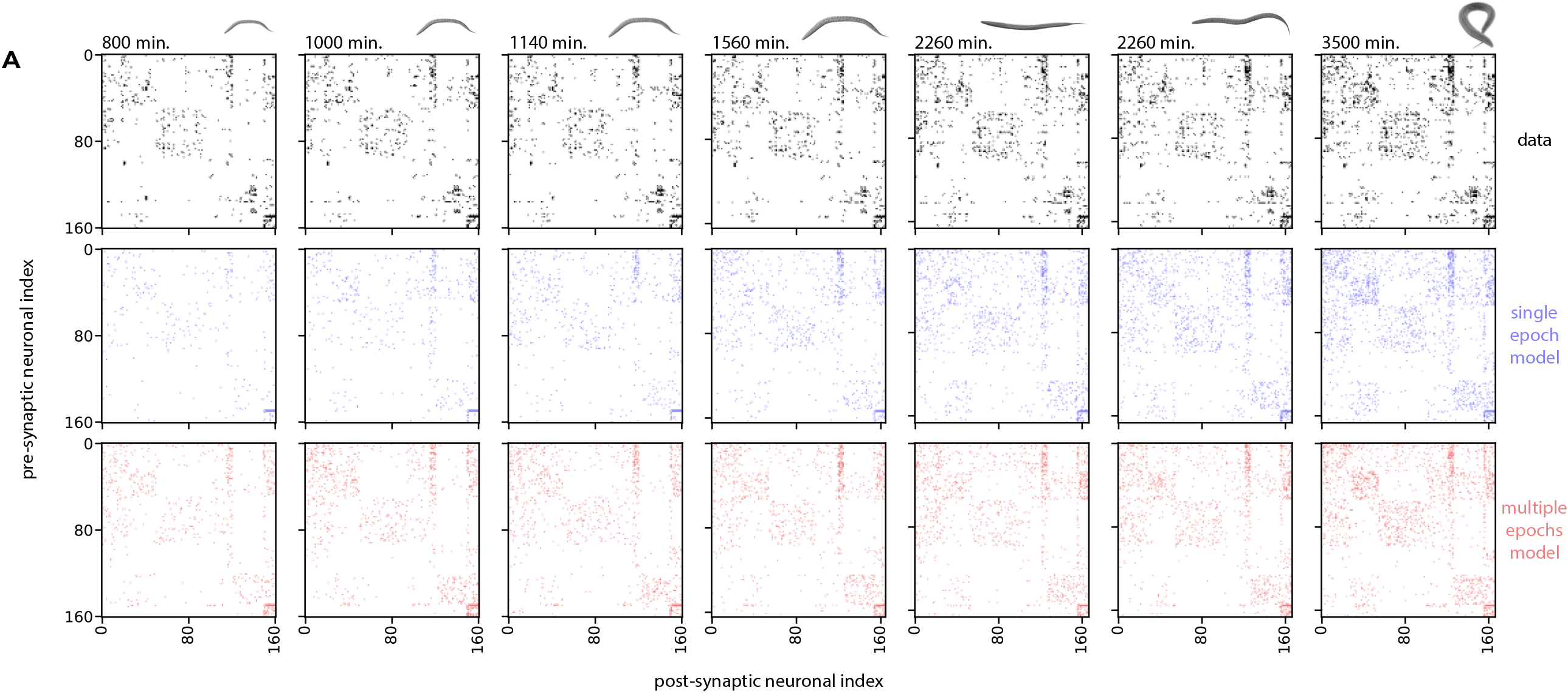
**A**. Connectivity maps of data (top) and of single draws from the single epoch model (middle) and the multiple epochs model (bottom) at ages corresponding to worms 1-6 and 8 (adult) in [16].

**Figure S 5.**
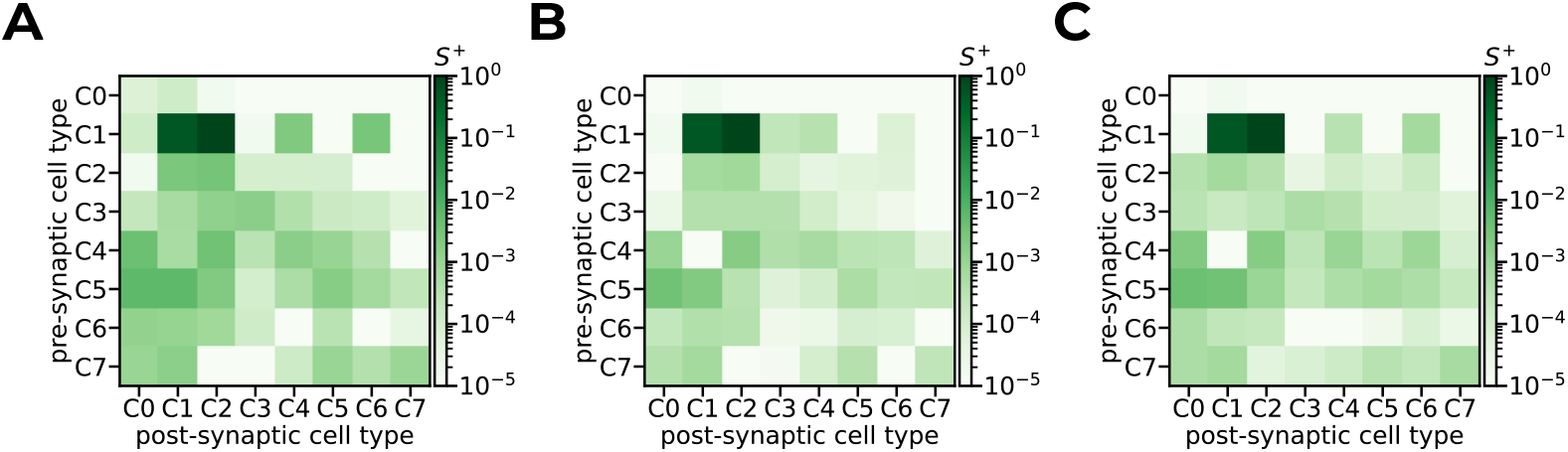
**A-C**. *S*^+^ matrices for the three developmental epochs (the order of the panels corresponds to the order of the epochs), for a single example split of all possible dyads into training and test subsets.

**Figure S 6.**
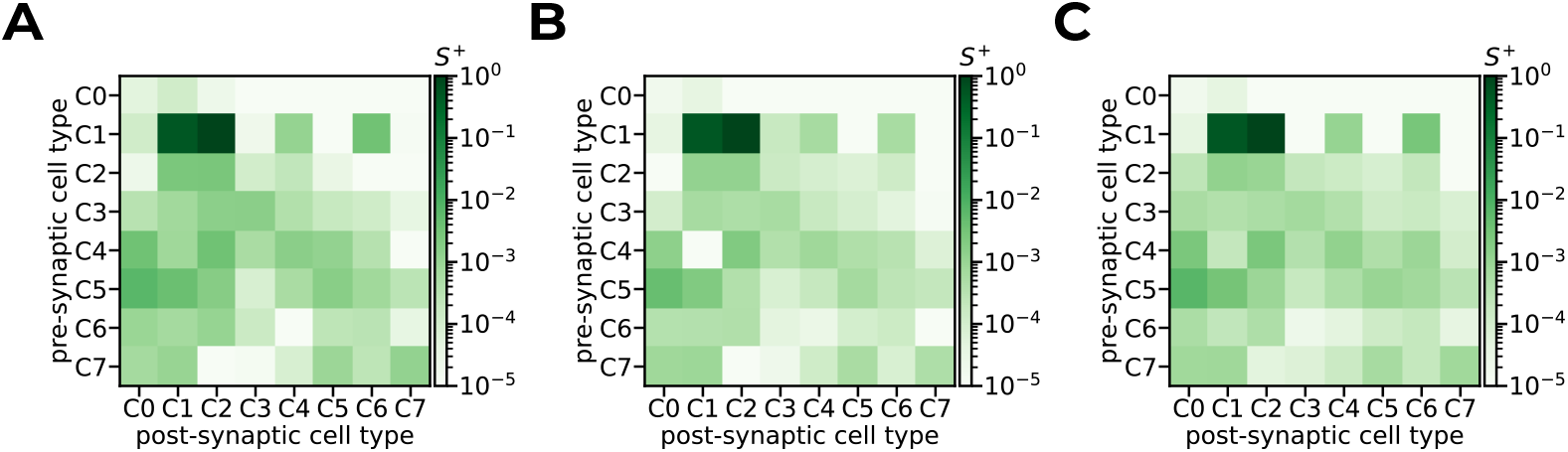
**A-C**. Average *S*^+^ matrix for the three developmental epochs (the order of the panels corresponds to the order of the epochs). Averages are over 20 splits of all possible dyads into training and test subsets.

**Figure S 7.**
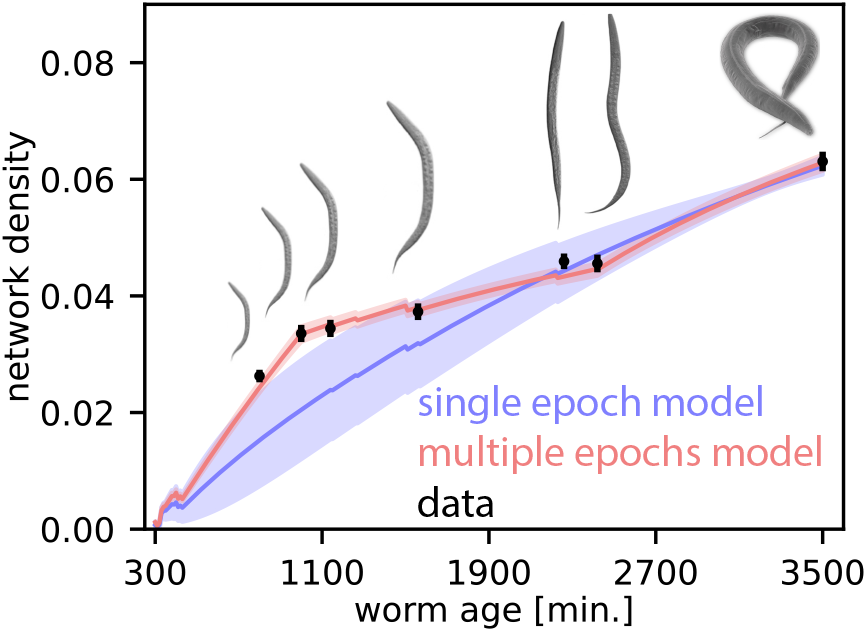
Multiple developmental epochs are necessary to follow the observed developmental path. Network density versus worm age, for the measured connectivity maps (test data), the single epoch model and the multiple epochs model. Solid lines represent models’ average density over 20 splits of the data into training and test subsets (the average of their mean densities), and the shaded areas are 1-std regions. Calculations are exact (see Methods). Error-bars represent the standard deviation of the density of the test subset across splits.

**Figure S 8.**
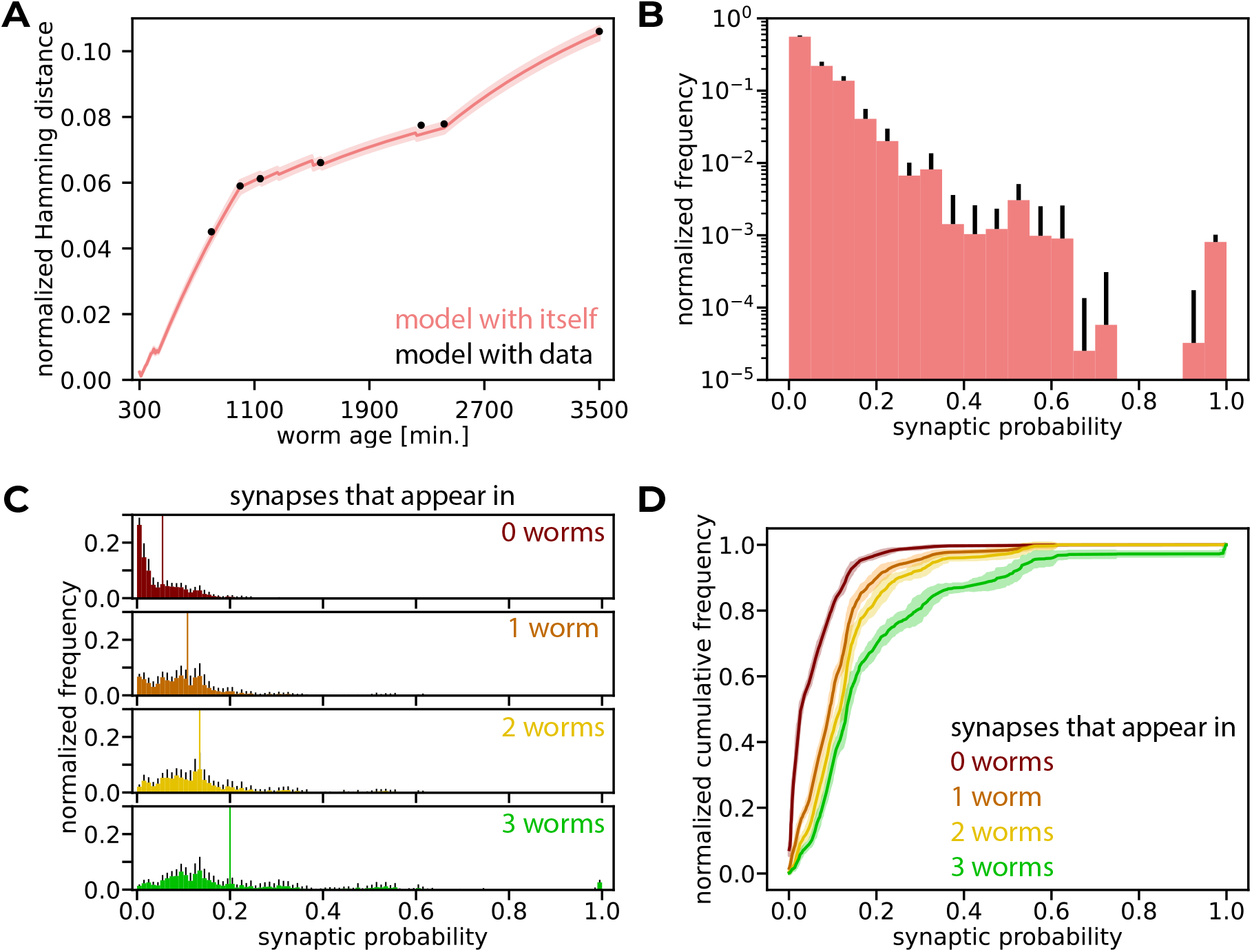
The model reveals the stable backbone of the connectivity map. **A**. The variance of the model and the cross-variance of the model and the data (measured by normalized Hamming distance). The solid line represents the average normalized Hamming distance between model runs over 20 splits into training and test data (the average of the mean normalized Hamming distance of models) and the shaded area is the 1-std region. The error-bars of the cross-variance of the model with the test data, which denote 1 std over 20 split into training and test, are smaller than the marker size. Calculations are exact (see Methods). **B**. The average normalized histogram of the probabilities that models assigned to all possible synapses over 20 splits into training and test data. Error-bars show the standard deviation. **C**. Average normalized histograms of synaptic probabilities predicted by the model over 20 splits into training and test data, when synapses are grouped by the number of datasets in which they appear. Error-bars show the standard deviation. Vertical lines show the average distributions’ means. **D**. Average normalized cumulative distributions of synaptic probabilities predicted by the model over 20 splits into training and test data, when synapses are grouped by the number of datasets in which they appear. Solid lines show the average cumulative distributions and the shaded areas 1-std ranges.

